# Nanopore direct RNA sequencing maps an Arabidopsis N6 methyladenosine epitranscriptome

**DOI:** 10.1101/706002

**Authors:** Matthew T. Parker, Katarzyna Knop, Anna V. Sherwood, Nicholas J. Schurch, Katarzyna Mackinnon, Peter D. Gould, Anthony Hall, Geoffrey J. Barton, Gordon G. Simpson

**Author notes:** Contributed equally (Matthew T. Parker, Katarzyna Knop, Anna V. Sherwood, Nicholas J. Schurch). Biomathematics and Statistics Scotland, The James Hutton Institute, Aberdeen, AB15 8QH, UK.

## Abstract

Understanding genome organization and gene regulation requires insight into RNA transcription, processing and modification. We adapted nanopore direct RNA sequencing to examine RNA from a wild-type accession of the model plant *Arabidopsis thaliana* and a mutant defective in mRNA methylation (m^6^A). Here we show that m^6^A can be mapped in full-length mRNAs transcriptome-wide and reveal the combinatorial diversity of cap-associated transcription start sites, splicing events, poly(A) site choice and poly(A) tail length. Loss of m^6^A from 3’ untranslated regions is associated with decreased relative transcript abundance and defective RNA 3′ end formation. A functional consequence of disrupted m^6^A is a lengthening of the circadian period. We conclude that nanopore direct RNA sequencing can reveal the complexity of mRNA processing and modification in full-length single molecule reads. These findings can refine Arabidopsis genome annotation. Further, applying this approach to less well-studied species could transform our understanding of what their genomes encode.

## Introduction

Patterns of pre-mRNA processing and base modifications determine eukaryotic mRNA coding potential and fate. Alternative transcripts produced from the same gene can differ in the position of the start site, the site of cleavage and polyadenylation, and the combination of exons spliced into the mature mRNA. Collectively termed the epitranscriptome, RNA modifications play crucial context-specific roles in gene expression^1, 2^. The most abundant internal modification of mRNA is methylation of adenosine at the N6 position (m^6^A). Knowledge of RNA modifications and processing combinations is essential to understand gene expression and what genomes really encode. RNA sequencing (RNAseq) is used to dissect transcriptome complexity: it involves copying RNA into complementary DNA (cDNA) with reverse transcriptases (RTs) and then sequencing the subsequent DNA copies. RNAseq reveals diverse features of transcriptomes, but limitations can include misidentification of 3′ ends through internal priming^3^, spurious antisense and splicing events produced by RT template switching^4, 5^, and the inability to detect all base modifications in the copying process^6^. The fragmentation of RNA prior to short-read sequencing makes it difficult to interpret the combination of authentic RNA processing events and remains an unsolved problem^7^.

We investigated whether long-read direct RNA sequencing (DRS) with nanopores^8^ could reveal the complexity of Arabidopsis mRNA processing and modifications. In nanopore DRS, the protein pore (nanopore) sits in a membrane through which an electrical current is passed, and intact RNA is fed through the nanopore by a motor protein^8^. Each RNA sequence within the nanopore (5 bases) can be identified by the magnitude of signal it produces. Arabidopsis is a pathfinder model in plant biology, and its genome annotation strongly influences the annotation and our understanding of what other plant genomes encode. We applied nanopore DRS and Illumina RNAseq to wild-type Arabidopsis (Col-0) and mutants defective in m^6^A^9^ and exosome-mediated RNA decay^10^. We reveal m^6^A and combinations of RNA processing events (alternative patterns of 5′ capped transcription start sites, splicing, 3′ polyadenylation and poly(A) tail length) in full-length Arabidopsis mRNAs transcriptome-wide.

## Results

### Nanopore DRS detects long, complex mRNAs and short, structured non-coding RNAs

We purified poly (A)+ RNA from four biological replicates of 14-day-old Arabidopsis Col-0 seedlings. We incorporated synthetic External RNA Controls Consortium (ERCC) RNA Spike-In mixes into all replicates^11, 12^ and carried out nanopore DRS. Illumina RNAseq was performed in parallel on similar material. Using Guppy base-calling (Oxford Nanopore Technologies) and minimap2 alignment software^13^, we identified around 1 million reads per sample (Supplementary table 1). The longest read alignments were 12.7 kb for mRNA transcribed from AT1G48090, spanning 63 exons, (Figure 1A), and 12.8 kb for mRNA transcribed from At1G67120, spanning 58 exons (Supplementary figure 1A). These represent some of the longest contiguous mRNAs sequenced from Arabidopsis. Among the shortest read alignments were those spanning genes encoding highly structured non-coding RNAs such as UsnRNAs and snoRNAs such as U3 (Figure 1B).

**Figure 1.**
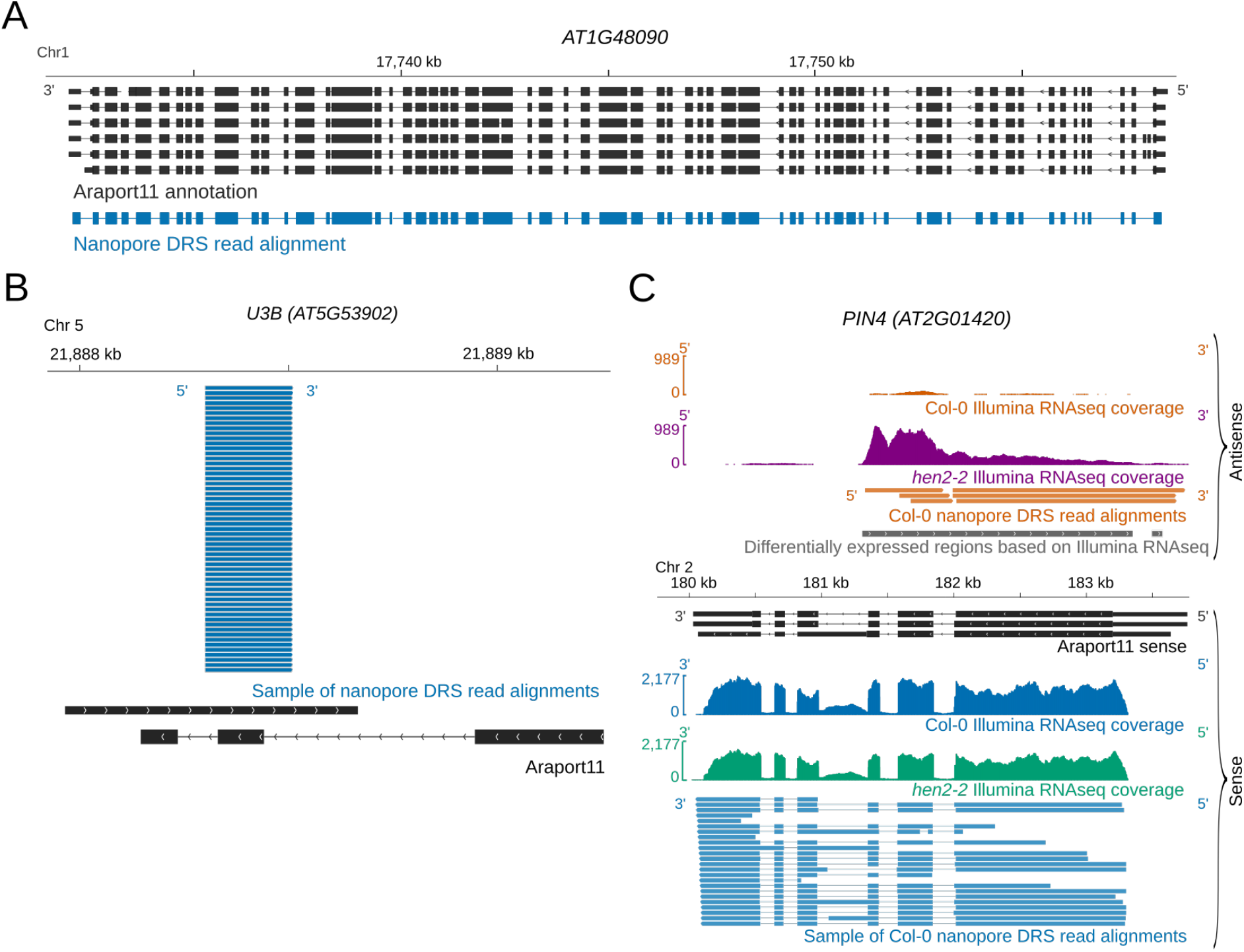
Diverse Arabidopsis RNAs are detected by nanopore DRS. (**A**) Nanopore DRS 12.7-kb read alignment at *AT1G48090*, comprising 63 exons. Black, Araport11 annotation; blue, nanopore DRS read alignment. (**B**) Nanopore DRS read alignments at the snoRNA gene *U3B*. Black, Araport11 annotation; blue, nanopore DRS read alignments. (**C**) *PIN4* long non-coding antisense RNAs detected using nanopore DRS. Blue, Col-0 sense Illumina RNAseq coverage and nanopore sense read alignments; orange, Col-0 antisense Illumina RNAseq coverage and nanopore antisense read alignments; green, *hen2-2* mutant sense Illumina RNAseq coverage; purple, *hen2-2* mutant antisense Illumina RNAseq coverage; black, sense RNA isoforms found in Araport11; grey, antisense differentially expressed regions detected with DERfinder. **[Linked to Supplementary figure 1].**

### Base-calling errors in nanopore DRS are non-random

We used ERCC RNA Spike-Ins^11^ as internal controls to monitor the properties of the sequencing reads. The spike-ins were detected in a quantitative manner (Supplementary figure 1B), consistent with the suggestion that nanopore sequencing is quantitative^8^. For the portion of reads that align to the reference, sequence identity was 92% when measured against the ERCC RNA spike-ins (Supplementary figure 1C). The errors showed evidence of base specificity (Supplementary figure 1D, E). For example, guanine was under-represented and uracil over-represented in indels and substitutions relative to the reference nucleotide (nt) distribution. In some situations, this bias could impact the utility of interpreting nanopore sequence errors. We used the proovread software tool^14^ and parallel Illumina RNAseq data to correct base-calling errors in the nanopore reads^15^.

### Artefactual splitting of raw signal affects transcript interpretation

We detected artefacts caused by the MinKNOW software splitting raw signal from single molecules into two or more reads. As a result, alignments comprising apparently novel 3′ ends were mapped as adjacent to alignments with apparently novel 5′ ends (Supplementary figure 1F). A related phenomenon called over-splitting was recently reported in nanopore DNA sequencing^16^. Over-splitting can be detected when two reads sequenced consecutively through the same pore are mapped to adjacent loci in the genome^16^. Over-splitting in nanopore DRS generally occurs at low frequency (< 2% of reads). However, RNAs originating from specific gene loci, such as *RH3* (*AT5G26742*), appear to be more susceptible, with up to 20% of reads affected across multiple sequencing experiments (Supplementary figure 1F).

### Spurious antisense reads are rare or absent in nanopore DRS

Since only two out of 9,445 (0.02%) reads mapped antisense to the ERCC RNA Spike-In collection^11^ and 0 of 19,665 reads mapped antisense to the highly expressed gene *RUBISCO ACTIVASE* (*RCA*) (*AT2G39730*), we conclude that spurious antisense is rare or absent from nanopore DRS data. This simplifies the interpretation of authentic antisense RNAs, which is important in Arabidopsis because the distinction between RT-dependent template switching and authentic antisense RNAs produced by RNA-dependent RNA polymerases that copy mRNA is not straightforward^17^. For example, by nanopore DRS, we could identify Arabidopsis long non-coding antisense RNAs, such as those at the auxin efflux carriers *PIN4* and *PIN7* (Figure 1C, Supplementary figure 1G). The existence of these previously unannotated antisense RNAs was supported by Illumina RNAseq of wild-type Col-0 and the exosome mutant *hen2–2* (Figure 1C, Supplementary figure 1G), the latter of which had a 13-fold increase in abundance of these antisense RNAs. Consequently, the low level of steady-state accumulation of some antisense RNAs may explain why they are currently unannotated.

### Nanopore DRS confirms sites of RNA 3′ end formation and estimates poly(A) tail length

Ligation of the motor protein adapter to RNA 3′ ends results in nanopores sequencing mRNA poly(A) tails first. We used the nanopolish-polyA software tool to estimate poly(A) tail lengths for individual transcripts^18^. This approach indicated an average length of 76 nt for Arabidopsis mRNA poly(A) tails, but with a wide range in estimated lengths for individual mRNAs (95% were in the 13–197 nt range; Figure 2A). The generally shorter poly(A) tails of chloroplast- and mitochondria-encoded transcripts, which are a feature of RNA decay in these organelles, were also detectable. We found that poly(A) tail length correlates negatively with gene expression in Arabidopsis (Spearman’s ρ=−0.3, *p*=2×10^-133^, 95% CIs [−0.32, −0.28]; Supplementary figure 2A), consistent with other species analysed by short-read TAILseq^19^.

**Figure 2.**
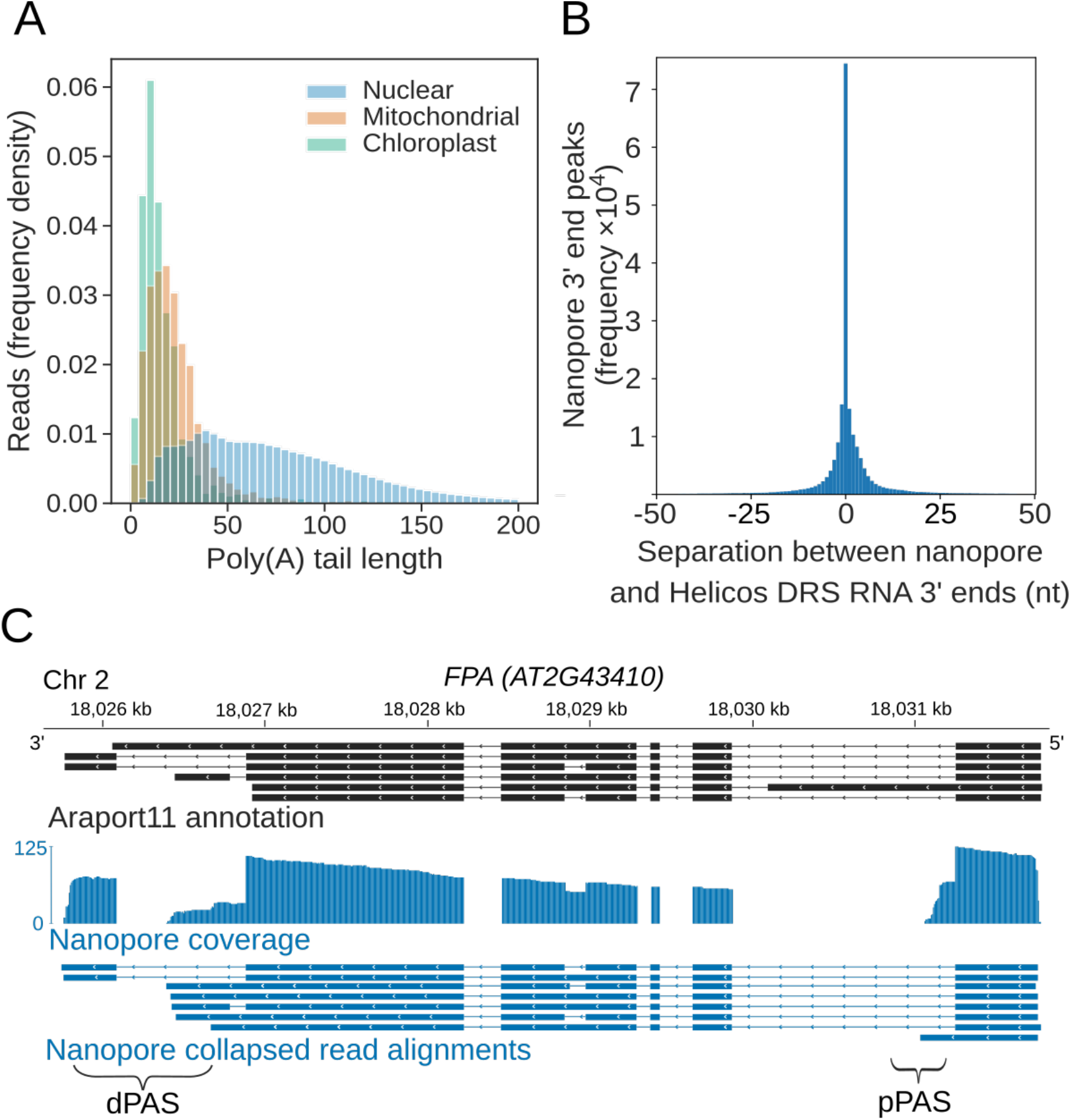
Nanopore DRS reveals poly(A) tail length and maps 3′ cleavage and polyadenylation sites. (**A**) Normalized histogram showing poly(A) tail length of RNAs encoded by different genomes. Blue, nuclear (n = 2,348,869 reads); orange, mitochondrial (n = 2,490 reads); green, chloroplast (n = 1,848 reads). (**B**) Distance between the RNA 3′ end positions in nanopore DRS read alignments and the nearest polyadenylation sites identified by Helicos data. (**C**) Nanopore DRS identified 3′ polyadenylation sites in RNAs transcribed from *FPA (AT2G43410)*. The blue track shows the coverage of nanopore DRS read alignments and collapsed read alignments representing putative transcript annotations detected by nanopore DRS. Black, isoforms found in Araport11 annotation; blue, read alignments from nanopore DRS. pPAS, proximal polyadenylation site; dPAS, distal polyadenylation sites. **[Linked to Supplementary figure 2].**

We previously mapped Arabidopsis mRNA 3′ ends transcriptome-wide using Helicos sequencing^20^. We compared the position of 3′ ends of nanopore DRS read alignments and Helicos data genome-wide. The median genomic distance between nanopore DRS and Helicos 3′ ends was 0 ± 13 nt (one standard deviation) demonstrating close agreement between these orthogonal technologies (Figure 2B). Likewise, the overall distribution of the 3′ ends of aligned nanopore DRS reads resembles the pattern we previously reported with Helicos data^20^. For example, 97% of nanopore DRS 3′ ends (4,152,800 reads at 639,178 unique sites, 93% of all unique sites) mapped to either annotated 3′ untranslated regions (UTRs) or downstream of the current annotation. Mapping of 3′ ends to coding sequences or 5′UTRs was rare (2.8%, 119,524 reads at 39,610 unique sites, 5.8% of all unique sites), and mapping to introns even rarer (0.29%, 12,554 reads at 7,791 unique sites, 1.1% of unique sites). Even so, examples of the latter included sites of alternative polyadenylation with well-established regulatory roles, such as in mRNA encoding the RNA-binding protein FPA, which controls flowering time^21^ (Figure 2C), and in mRNA encoding the histone H3K9 demethylase IBM1, which controls levels of genic DNA methylation^22^ (Supplementary figure 2B).

Since RT-dependent internal priming can result in the misinterpretation of authentic cleavage and polyadenylation sites^3^, we next determined whether nanopore DRS was compromised in this way. To address this issue, we examined whether the 3′ ends of nanopore DRS reads mapped to potential internal priming substrates comprised of six consecutive adenosines within a transcribed coding sequence (according to the Arabidopsis Information Portal Col-0 genome annotation, Araport11). Of the 10,116 such oligo (A)_6_ sequences, only four have read alignments terminating within 13 nt in all four datasets. Of these, two were not detectable after error correction with proovread (suggesting that they resulted from alignment errors) and the other two mapped to the terminal exon of coding sequence annotation, indicating that they may be authentic 3′ ends. Hence, internal priming is rare or absent in nanopore DRS data. Overall, we conclude that nanopore DRS can identify multiple authentic features of RNA 3′ end processing.

### Cap-dependent 5′ RNA detection by nanopore DRS

Nanopore DRS reads are frequently truncated prior to annotated transcription start sites, resulting in a 3′ bias of genomic alignments (Figure 3A)^15^. Consequently, it is impossible to determine which, if any, aligned reads correspond to full-length transcripts. To address this issue, we used cap-dependent ligation of a biotinylated 5′ adapter RNA to purify capped mRNAs. We then re-sequenced two biological replicates of Arabidopsis Col-0 incorporating 5′ adapter ligation (Supplementary table 1) and filtered the reads for 5′ adapter RNA sequences using the sequence alignment tool BLASTN and specific criteria (Supplementary table 2). We then used high confidence examples of sequences that passed or failed these criteria to train a convolutional neural network to detect the 5′ adapter RNA in the raw signal (Supplementary figure 3A–C). Hence, we improved 5′ adapter-ligated RNA detection without requiring base-calling or genome alignment, and demonstrated enrichment of full-length, cap-dependent mRNA sequences (Figure 3A, B). This procedure reduced the median 3′ bias of nanopore read alignments per gene (as measured by quartile coefficient of variation of per base coverage) from 0.45 (95% CIs [0.43,0.47]) to 0.08 (95% CIs [0.07,0.09]; Figure 3B).

**Figure 3.**
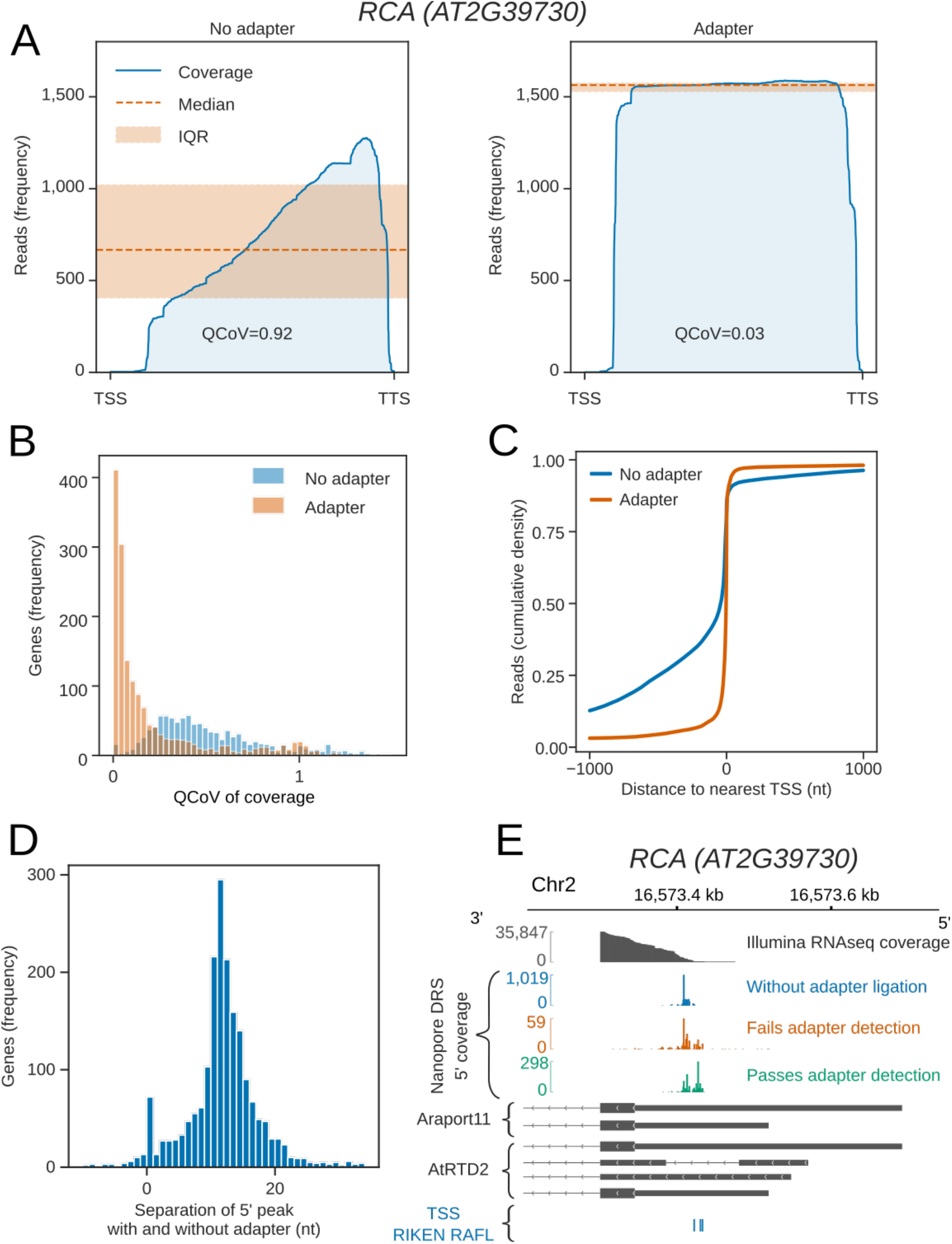
Cap-dependent ligation of an adapter enables detection of authentic RNA 5′ ends. (**A**) 5′ adapter RNA ligation reduces 3′ bias in nanopore DRS data at *RCA* (*AT2G39730*). Blue line, exonic read coverage at *RCA* for reads without (left) and with(right) adapter; orange line, median coverage; orange shaded area, interquartile range (IQR). Change in 3′ bias can be measured using the IQR / median = quartile coefficient of variation (QCoV). 5′ adapter ligation reduces 3′ bias at *RCA* from 0.92 to 0.03. (**B**) 5′ adapter RNA ligation reduces 3′ bias in nanopore DRS data. Histogram showing the QCoV in per base coverage for reads with a 5′ adapter RNA (orange) compared with reads without a 5′ adapter RNA (blue). (**C**) Cap-dependent adapter ligation allows identification of authentic 5′ ends using nanopore DRS. The cumulative distribution function shows the distance to the nearest Transcription Start Site (TSS) identified from full-length transcripts cloned as part of the RIKEN RAFL project for reads with a 5′ adapter RNA (orange) compared with reads without a 5′ adapter RNA (blue). (**D**) Cap-dependent adapter ligation enabled resolution of an additional 11 nt of sequence at the RNA 5′ end. Histogram showing the nucleotide shift in the largest peak of 5′ coverage for each gene in data obtained using protocols with vs without a 5′ adapter. (**E**) For *RCA (AT2G39730),* the 5′ end identified using cap-dependent 5′ adapter RNA ligation protocol was consistent with Illumina RNAseq and full-length cDNA start site data but differed from the 5′ ends in the Araport11 and AtRTD2 annotations. Upper panel: grey, Illumina RNAseq coverage; blue, nanopore DRS 5′ end coverage generated without a cap-dependent ligation protocol; green/orange, nanopore DRS 5′ end coverage for read alignments generated using the cap-dependent ligation protocol with (green) and without (orange) 5′ adapter RNA. Lower panel: grey, RNA isoforms found in Araport11 and AtRTD2 annotations; blue, TSSs identified from full-length transcripts cloned as part of the RIKEN RAFL project. **[Linked to Supplementary figure 3].**

In order to determine whether the 5′ ends we detected reflect full-length mRNAs, we compared them against annotated transcription start sites in datasets derived from full-length Arabidopsis cDNA clones^23^. We found that 41% of adapter-ligated nanopore DRS reads mapped within 5 nt of transcription start sites and 60% mapped within 13 nt (Figure 3C). We also detected recently defined examples of alternative 5′ transcription start sites^24^ at specific Arabidopsis genes (Supplementary figure 3D). We therefore conclude that this approach is effective in detecting authentic mRNA 5′ ends.

Reads with adapters had, on average, 11 nt more at their 5′ ends that could be aligned to the genome compared with the most common 5′ alignment position of reads lacking the 5′ adapter RNA (Figure 3D). This difference may be explained by loss of processive control by the motor protein when the end of an RNA molecule enters the pore. As a result, the 5′ end of RNA is not correctly sequenced. Consistent with these Arabidopsis transcriptome-wide nanopore DRS data, reads mapping to the synthetic ERCC RNA Spike-Ins and *in vitro* transcribed RNAs also lacked ∼11 nt of authentic 5′ sequence (Supplementary figure 3E, F). However, the precise length of 5′ sequence missing from all of these RNAs varied, suggesting that sequence- or context-specific effects on sequence accuracy are associated with the passage of 5′ RNA through the pore (Figure 3D, Supplementary figure 3E, F).

Despite the close agreement between nanopore DRS, Illumina RNAseq and full-length cDNA data^23^ at *RCA*, the start site annotated in Araport11 and the *Arabidopsis thaliana* Reference Transcript Dataset 2 (AtRTD2)^25^ is quite different (Figure 3E). The apparent overestimation of 5′UTR length is widespread in Araport11 annotation (Supplementary figure 3G), consistent with the assessment of capped Arabidopsis 5′ ends detected by nanoPARE sequencing^26^. Consequently, with appropriate modification to the current protocol, such as we describe here, nanopore DRS data can be used to revise Arabidopsis transcription start site annotations.

### Nanopore DRS reveals the complexity of splicing events

In single reads, nanopore sequencing revealed some of the most complex splicing combinations so far identified in the Arabidopsis transcriptome. For example, the splicing pattern of a 12.7 kb read alignment, comprised of 63 exons, agreed exactly with the *AT1G48090.4* isoform annotated in Araport11 (Figure 1A). Mutually exclusive alternative splicing of *FLM (AT1G77080*) exons that mediate the thermosensitive response controlling flowering time^27^ was also detected (Figure 4A). However, a combination of base-calling and alignment errors contributed to the misidentification of splicing events for uncorrected DRS data: 58% (170,702) of the unique splice junctions detected in the combined set of replicate data were absent from Araport11 and AtRTD2 annotations and were unsupported by Illumina RNAseq (Figure 4B, Supplementary table 3). We applied proovread^14, 15^ error correction with the parallel Illumina RNAseq data and then re-analysed the corrected and uncorrected nanopore DRS data. After error correction, only 13% (39,061) of unique splice junctions were unsupported by an orthogonal dataset, consistent with an improvement in alignment accuracy. The four nanopore DRS datasets for Col-0 biological replicates captured 75% (102,486) and 69% 104,686) of Araport11 and AtRTD2 splice site annotations, respectively. Most of the canonical GU/AG splicing events (100,450; 81%) detected in the error-corrected nanopore data were found in both annotations and were supported by Illumina RNAseq (Figure 4B, Supplementary table 3). A total of 3,234 unique canonical splicing events in the error-corrected nanopore DRS data were supported by Illumina RNAseq but absent from both Araport11 and AtRTD2 annotations, highlighting potential gaps in our understanding of the complexity of Arabidopsis splicing annotation (Figure 4B, Supplementary table 3). Consistent with this, we validated three of these splicing events using RT-PCR (Polymerase Chain Reaction) followed by cloning and sequencing (Supplementary figure 4A). In order to examine the features of these unannotated splices, we applied previously determined splice site position weight matrices of the flanking sequences to categorize U2 or U12 class splice sites^28^. Of the 3,234 novel GU/AG events found in error-corrected data and supported by Illumina alignments, 74% were classified as canonical U2 or U12 splice sites, suggesting that they are authentic (Supplementary figure 4B).

**Figure 4.**
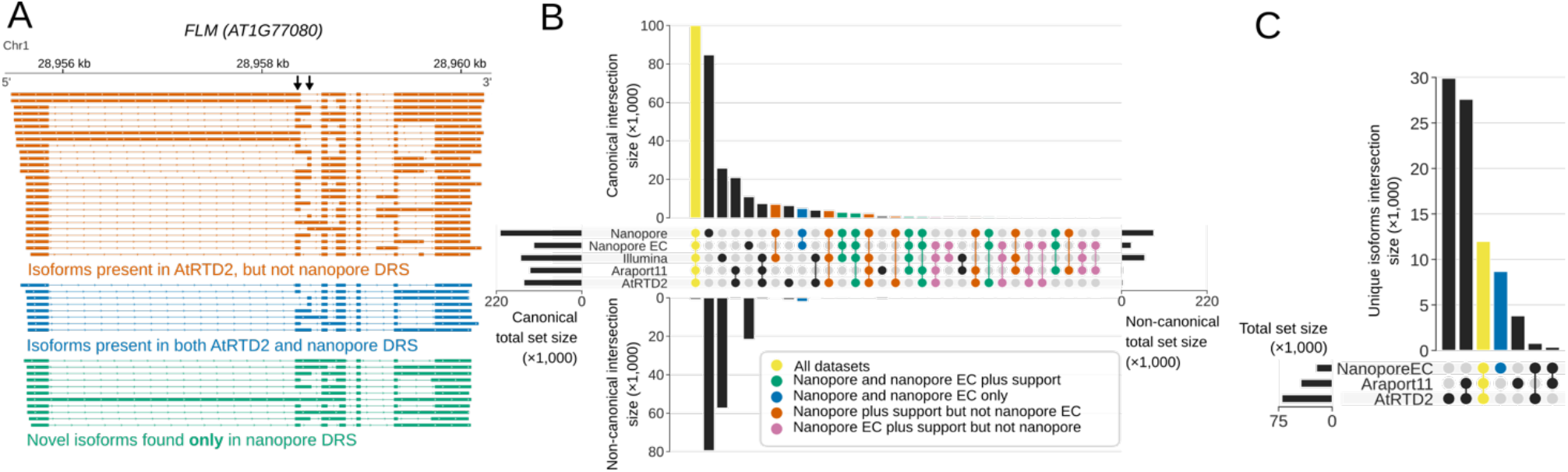
Nanopore DRS reveals the complexity of alternative splicing. (**A**) Nanopore DRS identified the mutually exclusive alternative splicing of *FLOWERING LOCUS M* (*FLM, AT1G77080)*. Black arrows indicate mutually exclusive exons. Novels isoforms were also identified: orange, isoforms present in the AtRTD2 annotation but not identified using nanopore DRS; blue, isoforms common to both AtRTD2 and nanopore DRS; green, novel isoforms identified in nanopore DRS. (**B**) Comparison of splicing events identified in error-corrected and non-error-corrected nanopore DRS, Illumina RNA sequencing, and Araport11 and AtRTD2 annotations. Bar size represents the number of unique splicing events common to the set intersection highlighted using circles (see Supplementary table 3 for the exact values). GU/AG splicing events are shown on the top and non-GU/AG on the bottom of the plot: yellow, splicing events common to all five datasets; green, events common to both error-corrected and non-error-corrected nanopore DRS with support in orthogonal datasets; blue, events common to both nanopore DRS datasets without orthogonal support; orange, events found in uncorrected nanopore DRS (but not error corrected) with orthogonal support; pink, events found in error-corrected nanopore DRS (but not uncorrected) with orthogonal support. (**C**) Comparison of RNA isoforms (defined as sets of co-spliced introns) identified in error-corrected full-length nanopore DRS, Araport11 and AtRTD2 annotations. Bar size represents the number of splicing events common to a group highlighted using circles below (see Supplementary table 3 for the exact values): yellow, unique splicing patterns nanopore DRS and both reference annotations; blue, novel isoforms. **[Linked to Supplementary figure 4].**

In addition to previously unannotated splicing events, we identified unannotated combinations of previously established splice sites. For example, we identified 19 *FLM* splicing patterns that adhered to known splice junction sites (Figure 4A). However, 11 of these transcript isoforms were not previously annotated. In order to investigate this phenomenon transcriptome-wide, we analysed the 5′ cap-dependent nanopore DRS datasets of full-length mRNAs (Supplementary table 1). Unique sets of co-splicing events were extracted from error-corrected reads (so as to focus on splicing, we did not consider single exon reads or 5′ and 3′ positions). In total, 13,064 unique splicing patterns were detected that matched annotations in Araport11, AtRTD2 or both (Figure 4C). Another 8,659 unique splicing patterns were identified that were not present in either annotation (Figure 4C, Supplementary table 3). Of these, 50% (4,293) used only splice donor and acceptor pairs that were already annotated in either Araport11 or AtRTD2. Hence, this approach defines splicing patterns (including retained introns) produced from alternative combinations of known splice sites.

Overall, we conclude that nanopore DRS can reveal a greater complexity of splicing in the context of full-length mRNAs compared with short-read data. However, accurate splice pattern detection benefits from error correction with, for example, high-accuracy orthogonal short-read sequencing data. However, even with error-free sequences, accurate splice detection can be confounded by the existence of equivalent alternative junctions^29^. Therefore, improved computational tools are required, not only for error correction but also for splicing-aware long-read alignment.

### Differential error site analysis reveals the m^6^A epitranscriptome

The epitranscriptome has emerged recently as a crucial, but relatively neglected, layer of gene regulation^1, 2^. m^6^A has been mapped transcriptome-wide using approaches based on antibodies that recognize this mark^6, 30^. However, in principle, m^6^A can be detected by nanopore DRS^8^. Since m^6^A is not included in the training data for nanopore base-calling software, we asked whether its incorrect interpretation could be used to identify Arabidopsis m^6^A transcriptome-wide. For this, we applied nanopore DRS to four biological replicates of an Arabidopsis mutant defective in the function of Virilizer (*vir-1*), a conserved m^6^A writer complex component, and four biological replicates of a line expressing VIR fused to Green Fluorescent Protein (GFP) that restores VIR activity in the *vir-1* mutant background^9^ (Supplementary figure 5A). In parallel, we sequenced a set of six biological replicates with Illumina RNAseq. We then used a G-test statistical analysis to determine whether there was a differential error profile in alignments at each reference base between the mutant (defective m^6^A) and VIR-complemented lines. We identified 17,491 sites with a more than two-fold higher error rate (compared with the TAIR10 reference base) in the VIR-complemented line with restored m^6^A (Figure 5A). No VIR-dependent error sites mapped to either chloroplast or mitochondrial-encoded RNAs. In all, 99.8% of the differential error sites mapped within Araport11 annotated protein-coding genes. Motif analysis of these error sites revealed the DRAYH sequence (D=G or U or A, R=G or A, Y=C or U, H=A or C or U; Figure 5B, E value=3.3×10^-191^), which closely resembles the established m^6^A target consensus^1, 2^. In addition, like the established location of m^6^A sites in mRNAs^1, 2^, the error sites were preferentially found in 3′UTRs (Figure 5C). Since approximately 5 nt contribute to the observed current at a given time point in nanopore sequencing^8^, the presence of a methylated adenosine could affect the accuracy of base-calling for the surrounding nucleotides. Consistent with this, we identified 4,749 sequences matching the motif discovered at error sites (False Discovery Rate [FDR] < 0.1; Supplementary figure 5B), with a median of two error sites per motif (95% CIs [1, 7]). Overall, these results agree with the established and conserved properties of authentic m^6^A sites^1, 2^, suggesting that differential error sites can be used to identify thousands of m^6^A modifications in nanopore DRS datasets.

**Figure 5.**
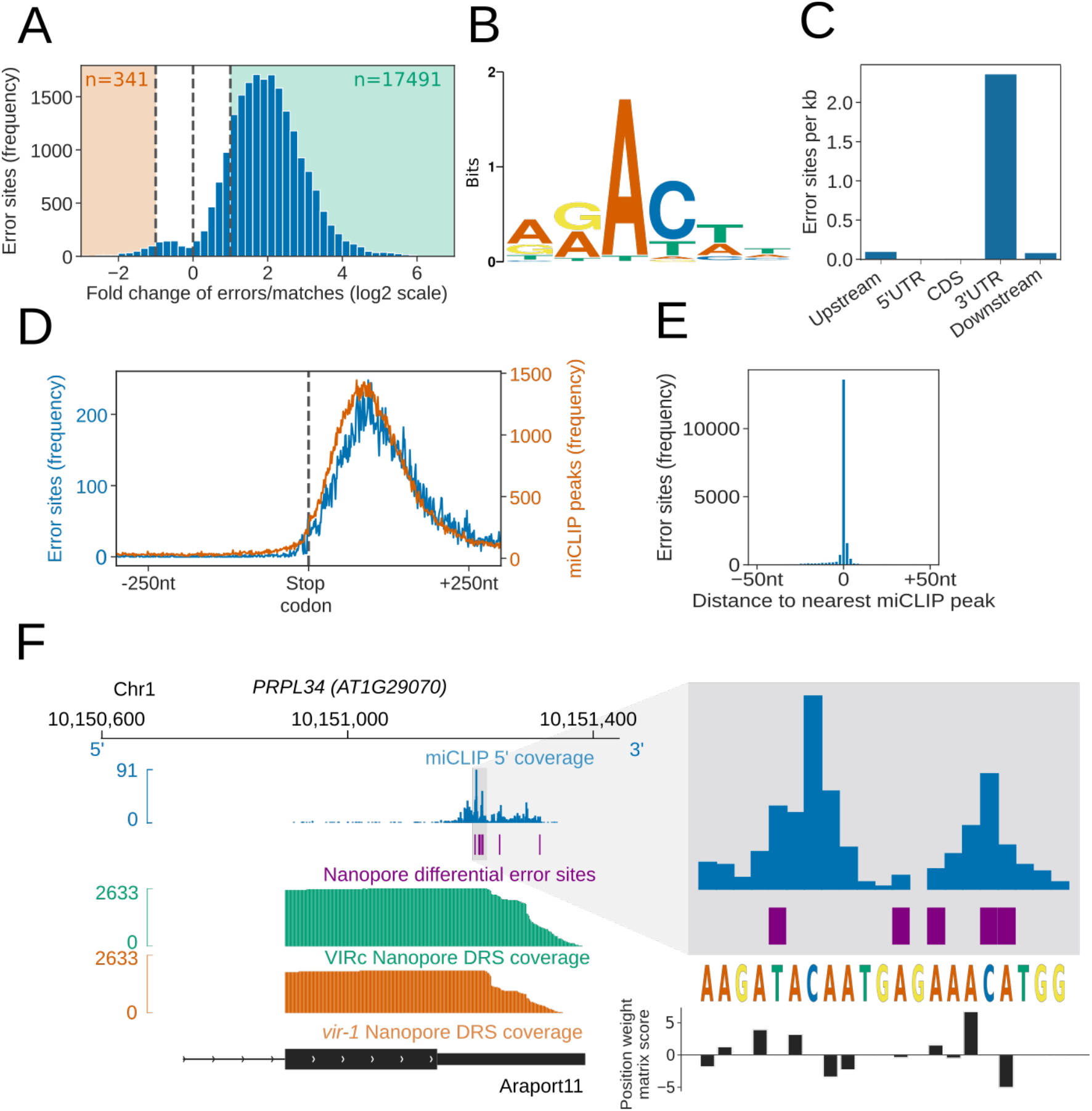
Differential error rate analysis identifies sites of VIR-dependent m^6^A modifications transcriptome-wide. (**A**) Loss of VIR function is associated with reduced error rate in nanopore DRS. Histogram showing the log2 fold change in the ratio of errors to reference matches at bases with a significant change in error profile in *vir-1* mutant compared with the VIR-complemented line. Orange and green shaded regions indicate sites with increased and reduced errors in *vir-1,* respectively. (**B**) The motif at error rate sites matches the consensus m^6^A target sequence. The sequence logo is for the motif enriched at sites with reduced error rate in the *vir-1* mutant. (**C**) Differential error rate sites are primarily found in 3′ UTRs. Bar plot showing the number of differential error rate sites per kb for different genic feature types of 48,149 protein coding transcript loci in the nuclear genome of the Araport11 reference. Upstream and downstream regions were defined as 200 nt regions 5′ and 3′ of the annotated transcription termination sites, respectively. (**D**) Differential error rate sites and miCLIP peaks are similarly distributed within the 3′ UTR, without accumulation at the stop codon. Metagene plot centred on stop codons from 48,149 protein coding transcript loci, showing the frequency of nanopore DRS error sites (blue) and miCLIP peaks (orange). (**E**) The locations of differential error rate sites are in good agreement with the locations of miCLIP sites. Histogram showing the distribution of distances to the nearest miCLIP peak for each site of reduced error. Most error sites (77%) are within 5 nt of a miCLIP peak. (**F**) Nanopore DRS differential error site analysis and miCLIP identify m^6^A sites in the 3′ UTR of *PRPL34* RNA. Blue, miCLIP 5′ end coverage; purple, nanopore DRS differential error sites; green, nanopore DRS coverage of VIR-complemented line (VIRc); orange, nanopore DRS coverage of *vir-1* mutant; black, RNA isoform from Araport11 annotation. The expanded region shows miCLIP coverage (blue) and error sites (purple) scored using the m^6^A consensus position weight matrix (black; Figure 5B). A higher positive score denotes a higher likelihood of a match to the consensus sequence. **[Linked to Supplementary figure 5].**

In order to examine the validity of m^6^A sites identified by the differential error site analysis, we used an orthogonal technique to map m^6^A. Previous maps of Arabidopsis m^6^A are based on Me-RIP^31, 32^ and limited by a resolution of around 200 nt^33^. Therefore, to examine Arabidopsis m^6^A sites with a more accurate method, we used miCLIP^30^ analysis of three biological replicates of Arabidopsis Col-0. We found that, like the differential error sites uncovered in the nanopore DRS analysis, the Arabidopsis miCLIP reads were enriched in 3′ UTRs but with no enrichment over stop codons (Figure 5D, Supplementary figure 5C). In all, 77% of the called nanopore DRS differential error sites fell within 5 nt of an miCLIP peak (Figure 5E, F). We therefore conclude that our analysis of nanopore data can detect authentic VIR-dependent m^6^A sites transcriptome-wide.

### Defective m^6^A perturbs gene expression patterns and lengthens the circadian period

The combination of transcript processing and modification data obtained using nanopore DRS enabled us to investigate the impact of m^6^A on Arabidopsis gene expression. We found a global reduction in protein-coding gene expression in *vir-1* (using either nanopore DRS or Illumina RNAseq data), corresponding to transcripts that were methylated in the VIR-complemented line (Figure 6A, Supplementary Figure 6A). These findings are consistent with the recent discovery that m^6^A predominantly protects Arabidopsis mRNAs from endonucleolytic cleavage^32^. Therefore, although the m^6^A writer complex comprises conserved components that target a conserved consensus sequence and distribution of m^6^A is enriched in the last exon, it appears that this modification predominantly promotes mRNA decay in human cells^34^, but mRNA stability in Arabidopsis^32^.

**Figure 6.**
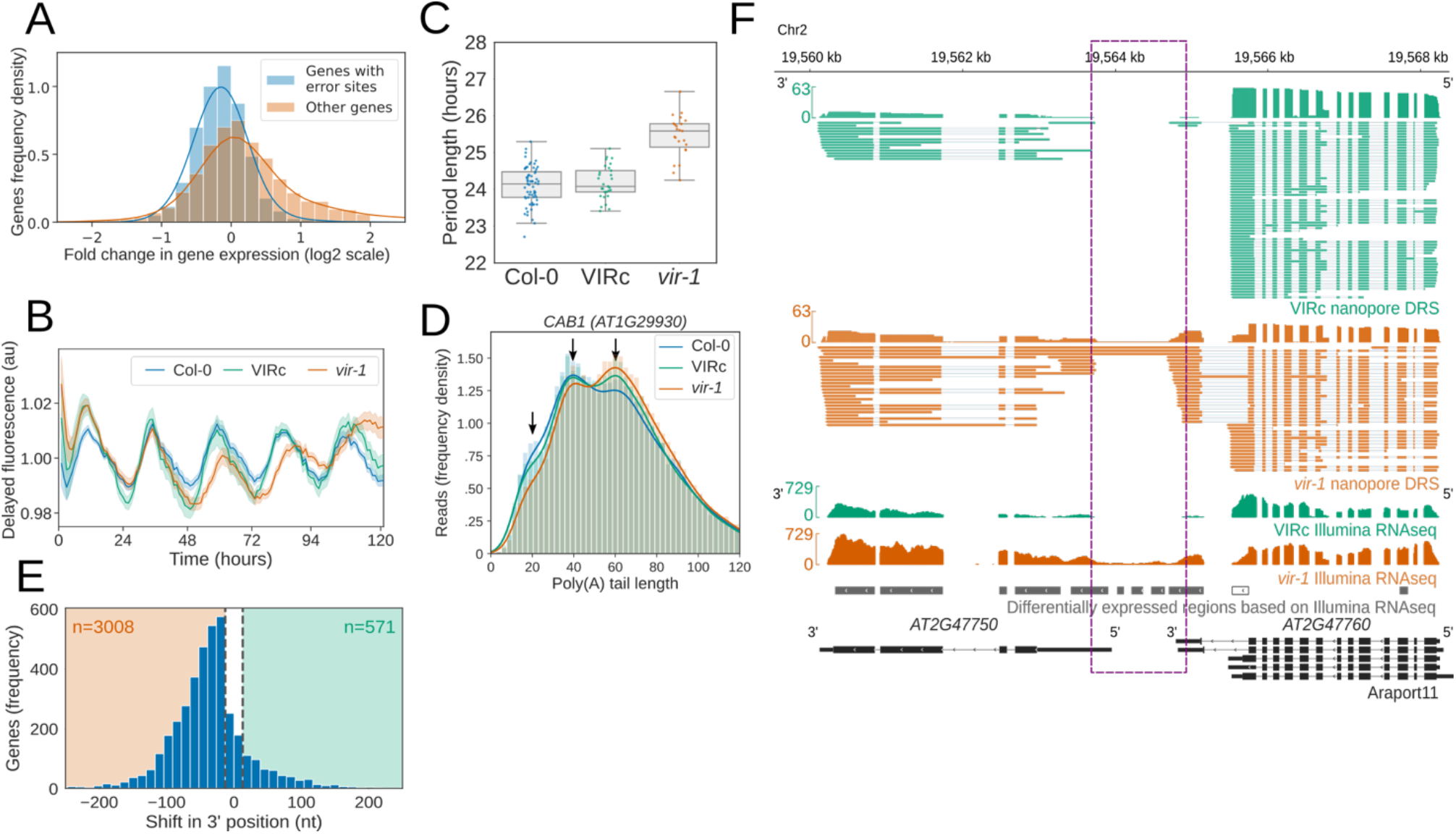
Reduction in m^6^A RNA modification leads to disruption of the circadian clock and generation of chimeric RNAs. (**A**) Genes with differential error rate sites have lower detectable RNA levels. Histogram showing the log2 fold change in protein coding gene expression based on counts from nanopore DRS reads in the *vir-1* mutant compared to the VIR-complemented line. Blue, genes with differential error rate sites (n = 5,157 genes); orange, genes with without differential error rate sites (n=14,601 genes). (**B**) The circadian period is lengthened in the *vir-1* mutant. Mean delayed fluorescence measurements in arbitrary units are shown for Col-0 (blue; n = 61 technical reps), the VIR-complemented line (VIRc; green; n = 29 technical reps) and the *vir-1* mutant (orange; n = 24 technical reps). Shaded areas show bootstrapped 95% confidence intervals for the mean. (**C**) Boxplot showing the period lengths for Col-0 (blue), VIRc (green) and the *vir-1* mutant (orange), calculated from delayed fluorescence measurements shown in (**B**). (**D**) Poly(A) tail length is altered in the *vir-1* mutant. Histogram showing the poly(A) tail length distribution of *CAB1 (AT1G29930)* in Col-0 (blue; n=40,841 reads), VIRc (green; n=65,810 reads) and the *vir-1* mutant (orange; n=68,068 reads). Arrows indicate phased peaks of poly(A) length at approximately 20, 40 and 60 nt. *vir-1* distribution is significantly different from both Col-0 and VIRc, according to the Kolmogorov–Smirnov test (*p*=1.3×10^-76^, *p*=4.7×10^-49^ respectively). (**E**) Global change in 3′ end usage in the *vir-1* mutant compared with the VIRc line. Histogram showing the distance in nucleotides between the most reduced and most increased 3′ end positions for genes in which the 3′ end profile is altered in *vir-1* (detected with the Kolmogorov–Smirnov test, FDR < 0.05). A threshold of 13 nt was used to detect changes in the 3′ end position. (**F**) Readthough events and chimeric RNAs are detected in *vir-1*. Green, nanopore DRS and Illumina RNAseq data for the VIRc line; orange, nanopore DRS and Illumina RNAseq data for the *vir-1* mutant; black, RNA isoforms found in Araport11 annotation. Differentially expressed regions between *vir-1* and VIRc detected using Illumina RNAseq data with DERfinder are shown in grey (for upregulated regions) or white (for downregulated regions). Intergenic readthrough regions are highlighted by the purple dashed rectangle. **[Linked to Supplementary figure 6].**

The changes in gene expression in *vir-1* were wide ranging (Supplementary figure 6B–D). For example, we found that the abundance of mRNAs encoding components of the interlocking transcriptional feedback loops that comprise the Arabidopsis circadian oscillator^35^, such as *CCA1*, were altered in *vir-1* (Supplementary figure 6B, C). This distinction was associated with a biological consequence in that the *vir-1* mutant had a lengthened clock period (Figure 6B, C). We detected m^6^A at mRNAs encoding the clock components *CCA1*, *PRR7, GI* and *LNK1/2* in both the nanopore DRS and miCLIP data (Supplementary figure 6B). We also detected shifts in the poly(A) tail length distributions of mRNAs transcribed from genes previously shown to be under circadian control. At *CAB1* mRNAs, for example, we detected poly(A) tail lengths that peaked at approximately 20, 40 and 60 nt (Figure 6D). *vir-1* mutants had reduced abundance of *CAB1* mRNAs with 20 and 40 nt poly(A) tails, and an increased abundance of poly(A) tails of 60 nt in length (Figure 6D). Therefore, nanopore DRS may uncover the circadian control of poly(A) tail length, as previously reported for specific Arabidopsis genes^36^. An output of the circadian clock is the control of flowering time, and we found that not only were photoperiod pathway components differentially expressed but so too were other flowering time genes (Supplementary table 4). Notably, *FLOWERING LOCUS C* (*FLC*) expression was reduced by more than 40-fold compared with the wild type (Supplementary figure 6D). Consequently, the proper control of circadian rhythms, flowering time and the regulatory module at *FLC* ultimately requires the m^6^A writer complex component, VIR.

### Defective m^6^A is associated with disrupted RNA 3′ end formation

In addition to measuring RNA expression, we examined the impact of m^6^A loss on pre-mRNA processing. Detectable disruptions to splicing in *vir-1* were modest. For example, using the DEX-Seq software tool^37^ for analysis of annotated splice sites, we found only weak effect-size changes to cassette exons, retained introns or alternative donor/acceptor sites compared with the VIR-complemented line in our Illumina data (Supplementary figure 6E). In contrast, a clear defect in RNA 3′ end formation in *vir-1* was apparent. Using a Kolmogorov–Smirnov test, we identified 3,579 genes with an altered nanopore DRS 3′ position profile in the *vir-1* mutant compared with the VIR-complemented line (FDR < 0.05, absolute change in position of >13 nt; Figure 6E). Of these, 3,008 displayed a shift to usage of more proximal poly(A) sites in *vir-1*: 60% of these genes also contain m^6^A sites detectable by nanopore DRS (compared with 32% of all expressed genes, *p*=1.1×10^-266^; 70% were detectable by miCLIP compared with 43% of all expressed genes, *p*=3.5×10^-237^) and correspond to locations of increased cleavage downstream of m^6^A sites in the *vir-1* mutant (Supplementary figure 6F). A total of 571 genes showed increased transcriptional readthrough beyond the 3′ end in *vir-1* (Figure 6E): 73% of these loci also contained nanopore-mapped m^6^A sites (*p*=1.2×10^-90^; 78% were detectable by miCLIP, *p*=2.1×10^-66^). The impacts of altered 3′ processing can be complex and have the potential to change the relative abundance of transcripts processed from the same gene but with different coding potential. For example, we detected increased readthrough of an intronic cleavage site in the Symplekin-like gene *TANG1* (*AT1G27595*; Supplementary figure 6G) and increased readthrough and cryptic splicing at *ALG3* (*AT2G47760*) that also results in chimeric RNA formation with the downstream gene *GH3.9* (*AT2G4G7750*; Figure 6F). The existence of the *ALG3-GH3.9* chimeric RNAs was supported by Illumina RNAseq (Figure 6F) and confirmed by RT-PCR, cloning and sequencing (Supplementary figure 6H). We detected 523 loci with increased levels of chimeric RNAs in *vir-1* resulting from unterminated transcription proceeding into downstream genes on the same strand. Chimeric RNAs were recently detected in mutants affecting other components of the Arabidopsis m^6^A writer complex, MTA and FIP37^38^. However, only 33% of upstream genes forming the chimeric RNAs had detectable m^6^A sites in the VIR-complemented line with restored VIR activity. Consequently, these findings might be explained either by an m^6^A-independent role for VIR (or the writer complex) in 3′ end formation or an indirect effect on factors required for 3′ processing. m^6^A independent roles for the human m^6^A methyltransferases METTL3^39^ and METTL16^40^ have been found previously, and a role for the writer complex in controlling Arabidopsis RNA processing independent of m^6^A cannot be overlooked^40^. In mammals, recognition of the canonical poly(A) signal AAUAAA involves direct binding by CPSF30^41, 42^. Notably, alternative polyadenylation controls the expression of an Arabidopsis CPSF30 isoform that encodes a YT521-B homology (YTH) domain with the potential to bind and read m^6^A^43^. A recent study indicated that this YTH domain-containing isoform is required to supress chimera formation^38^. Consequently, in plants, m^6^A may also contribute to the recognition of specific RNA 3′ ends.

## Concluding remarks

We have shown that nanopore DRS has the potential to refine multiple features of Arabidopsis genome annotation and to generate the clearest map to date of m^6^A locations, despite the genome sequence being available since 2000^44^. Modern agriculture is dominated by a handful of intensely researched crops^45^, but to diversify global food supply, enhance agricultural productivity and tackle malnutrition there is a need to focus on crops utilized in rural societies that have received little attention for crop improvement^46^. Based on our experience with Arabidopsis, we anticipate that the combination of nanopore DRS and other sequencing approaches will improve genome annotation. Consistent with this, we recently applied nanopore DRS to refine the annotation of water yam (*Dioscorea alata*), an African orphan crop. Indeed, we are moving into an era where thousands of genome sequences are available and programmes such as the Earth BioGenome Project aim to sequence all eukaryotic life on Earth^47^. However, genome sequences provide only part of the puzzle: annotating what they encode will be essential for us to fully utilize this information.

## Materials and methods

### Plants

#### Plant material and growth conditions

Wild-type *A. thaliana* accession Col-0 was obtained from Nottingham Arabidopsis Stock Centre. The *vir-1* and VIR-complemented (*VIR::GFP-VIR*) lines were provided by K. Růžička, Brno^48^. The *hen2–2* (Gabi_774HO7) mutant was provided by D. Gagliardi, Strasbourg. Seeds were sown on MS10 medium plates, stratified at 4°C for 2 days, germinated in a controlled environment at 22°C under 16-h light/8-h dark conditions and harvested 14 days after transfer to 22°C.

#### Clock phenotype analysis

Clock phenotype experiments were performed as previously described by measuring delayed fluorescence as a circadian output^49^. Briefly, plants were grown in 12-h light/12-h dark cycles at 22°C and 80 μmol m^−2^ sec^−1^ light for 9 days. Next, delayed fluorescence measurements were recorded every hour for 6 days at constant temperature (22°C) and constant light (20 µmol m^−2^ sec^−1^ red light and 20 µmol m^−2^ sec^−1^ blue light mix). Fast Fourier Transform (FFT) non-linear least-squares fitting to estimate period length was conducted using Biodare^50^.

### RNA

#### RNA isolation

Total RNA was isolated using RNeasy Plant Mini kit (Qiagen) and treated with TURBO DNase (Thermo Fisher Scientific). The total RNA concentration was measured using a Qubit 1.0 Fluorometer and Qubit RNA BR Assay Kit (Thermo Fisher Scientific), and RNA quality and integrity were assessed using a NanoDrop 2000 spectrophotometer (Thermo Fisher Scientific) and Agilent 2200 TapeStation System (Agilent).

#### mGFP in vitro transcription

The mGFP coding sequence was amplified using CloneAmp HiFi PCR Premix (Clontech) and a forward primer containing the T7 promoter sequence (Merck; Supplementary table 5). The PCR product was purified using GeneJET Gel Extraction (Thermo Fisher Scientific) and DNA Cleanup Micro (Thermo Fisher Scientific) kits, according to the manufacturer’s instructions. mGFP was *in vitro* transcribed using a mMESSAGE mMACHINE T7 ULTRA Transcription Kit (Thermo Fisher Scientific) with and without the anti-reverse cap analogue, according to the manufacturer’s instructions. mGFP transcripts were treated with TURBO DNase, polyA-tailed using *Escherichia coli* poly(A) Polymerase (E-PAP) and ATP (Thermo Fisher Scientific), and recovered using a MEGAclear kit (Thermo Fisher Scientific) according to the manufacturer’s instructions. The quantity of mGFP mRNAs was measured using a Qubit 1.0 Fluorometer (as described above), and the quality and integrity was checked using the NanoDrop 2000 spectrophotometer and agarose-gel electrophoresis. *In vitro* capped and non-capped mGFP mRNAs were used to prepare the library for DRS using nanopores.

#### RT-PCR and RT-qPCR

Total RNA was reverse transcribed using SuperScript III polymerase or SuperScript IV VILO Master Mix (Thermo Fisher Scientific) according to the manufacturer’s protocol. For RT-PCR, reactions were performed using the Advantage 2 Polymerase Mix (Clontech) using the primers (Merck) listed in Supplementary table 5. PCR products were gel purified using GeneJET Gel Extraction and DNA Cleanup Micro kits (Thermo Fisher Scientific), cloned into the pGEM T-Easy vector (Promega; according to the manufacturer’s instruction) and sequenced. For RT-qPCR, reactions were carried out using the SYBR Green I (Qiagen) mix with the primers (Merck) listed in Supplementary table 5, following manufacturer’s instructions.

### Illumina RNA sequencing

#### Preparation of libraries for Illumina RNA sequencing

Illumina RNA sequencing libraries from purified mRNA were prepared and sequenced by the Centre for Genomic Research at University of Liverpool using the NEBNext Ultra Directional RNA Library Prep Kit for Illumina (New England Biolabs). Paired-end sequencing with a read length of 150 bp was carried out on an Illumina HiSeq 4000. Illumina RNA libraries from ribosome-depleted RNA were prepared using the TruSeq Stranded Total RNA with Ribo-Zero Plant kit (Illumina). Paired-end sequencing with a read length of 100 bp was carried out on an Illumina HiSeq 2000 at the Genomic Sequencing Unit of the University of Dundee. ERCC RNA Spike-In mixes (Thermo Fisher Scientific)^11, 51^ were included in each of the libraries using concentrations advised by the manufacturer.

#### Mapping of Illumina RNA sequencing data

Reads were mapped to the TAIR10 reference genome with Araport11 reference annotation using STAR version 2.6.1^52^, a maximum multimapping rate of 5, a minimum splice junction overhang of 8 nt (3 nt for junctions in the Araport11 reference), a maximum of five mismatches per read and intron length boundaries of 60–10,000 nt.

#### Differential gene expression analysis using Illumina RNA sequencing data

Transcript-level counts for Illumina RNA sequencing reads were estimated by pseudoalignment with Salmon version 0.11.2^53^. Counts were aggregated to gene level using tximport^54^ and differential gene expression analyses for *vir-1* mutant vs wild type and *vir-1* mutant vs the VIR-complemented line were conducted in R version 3.5 using edgeR version 3.24.3^55^.

#### Differentially expressed region analysis using Illumina RNA sequencing data

Mapped read pairs originating from the forward and reverse strands were separated and coverage tracks were generated using samtools version 1.9^56^. Coverage tracks were then used as input for DERfinder version 1.16.1^57^. Expressed regions were identified using a minimum coverage of 10 reads, and differential expression between the *vir-1* and VIR-complemented lines was assessed using the analyseChr method with 50 permutations.

#### Differential exon usage analysis using Illumina RNA sequencing data

Annotated gene models from Araport11 were divided into transcript chunks (i.e. contiguous regions within which each base is present in the same set of transcript models). Read counts for each chunk were generated using bedtools version 2.27.1^58^ intersect in count mode. Chunk counts were then processed using DEXseq version 1.28.3^37^ to identify differentially expressed chunks between *vir-1* and VIR-complemented lines, using an absolute log-fold-change threshold of 1 and an FDR threshold of 0.05. Chunks were annotated as 5′ variation if they included a start site of any transcript and as 3′ variation if they contained a termination site. Chunks representing overhangs from alternative donor or acceptor sites were also classified separately. Internal exons were subclassified as a cassette exon if they could be wholly contained within any intron.

### Nanopore DRS

#### Preparation of libraries for direct RNA sequencing using nanopores

mRNA was isolated from approximately 75 μg of total RNA using the Dynabeads mRNA purification kit (Thermo Fisher Scientific) following the manufacturer’s instructions. The quality and quantity of mRNA was assessed using the NanoDrop 2000 spectrophotometer (Thermo Fisher Scientific). Nanopore libraries were prepared from 1 μg poly(A)+ RNA combined with 1 μl undiluted ERCC RNA Spike-In mix (Thermo Fisher Scientific) using the nanopore DRS Kit (SQK-RNA001, Oxford Nanopore Technologies) according to manufacturer’s instructions. The poly(T) adapter was ligated to the mRNA using T4 DNA ligase (New England Biolabs) in the Quick Ligase reaction buffer (New England Biolabs) for 15 min at room temperature. First-strand cDNA was synthesized by SuperScript III Reverse Transcriptase (Thermo Fisher Scientific) using the oligo(dT) adapter. The RNA–cDNA hybrid was then purified using Agencourt RNAClean XP magnetic beads (Beckman Coulter). The sequencing adapter was ligated to the mRNA using T4 DNA ligase (New England Biolabs) in the Quick Ligase reaction buffer (New England Biolabs) for 15 min at room temperature followed by a second purification step using Agencourt beads (as described above). Libraries were loaded onto R9.4 SpotON Flow Cells (Oxford Nanopore Technologies) and sequenced using a 48-h run time.

To incorporate cap-dependent ligation of a biotinylated 5′ adapter RNA, the following modifications were introduced into the library preparation protocol. First, 4 µg mRNA was de-phosphorylated by calf intestinal alkaline phosphatase (Thermo Fisher Scientific) and the 5′ cap was removed by Cap-Clip Acid Pyrophosphatase (Cambio) according to the manufacturer’s instructions. Next, the 5′ RNA oligo biotinylated at the 5′ end (Integrated DNA Technologies) was ligated to dephosphorylated, de-capped mRNA using T4 RNA ligase I (New England Biolabs) and mRNA was purified using Dynabeads MyOne Streptavidin C1 beads (Thermo Fisher Scientific) according to the manufacturer’s instructions. mRNA was assessed for quality and quantity using the NanoDrop 2000 spectrophotometer and used for nanopore DRS library preparation (as described above).

#### Processing of nanopore DRS data

Reads were basecalled with Guppy version 2.3.1 (Oxford Nanopore Technologies) using default RNA parameters and converted from RNA to DNA fastq using seqkit version 0.10.0^59^. Reads were aligned to the TAIR10 *A. thaliana* genome^60^ and ERCC RNA Spike-In sequences^11, 51^ using minimap2 version 2.8^13^ in spliced mapping mode using a kmer size of 14 and a maximum intron size of 10,000 nt. Sequence Alignment/Map (SAM) and BAM file manipulations were performed using samtools version 1.9^56^.

Proovread version 2.14.1^14^ was used to correct errors in the nanopore DRS reads^15^. Each nanopore DRS replicate was split into 200 chunks for parallel processing. Each chunk was corrected using four samples of Illumina poly(A) RNAseq data selected randomly from the 36 Illumina files (six biological replicates sequenced across six lanes). Illumina reads 1 and 2 were merged into fragments using FLASH version 1.2.11^61^. Unjoined pairs were discarded. Error correction with proovread was conducted in sampling-free mode using a minimum nanopore read length of 50 nt. Since both the Illumina RNAseq and nanopore DRS datasets were strand specific, proovread was modified to prevent opposite strand mapping between the datasets. Corrected reads were then mapped to the Araport11 reference genome using minimap2 (as described above). All figures showing gene tracks with nanopore DRS reads use error-corrected reads.

#### Error profile analysis using nanopore DRS data

Error rate analysis of aligned reads was conducted on ERCC RNA Spike-In mix controls using pysam version 0.15.2^62^ for BAM file parsing. Matches, mismatches, insertions and deletions relative to the reference were extracted from the cs tag (a more informative version of CIGAR string, output by minimap2) and normalised by the aligned length of the read. Reference bases and mismatch bases per position were also recorded and used to assess the frequency of each substitution and indel type by reference base.

#### Over-splitting analysis of nanopore DRS data

To identify read pairs resulting from over-splitting of the signal originating from a single RNA molecule, the sequencing summary files produced by Guppy were parsed for sequencing time and channel identifier and then used to identify pairs of consecutively sequenced reads. Genomic locations of reads were parsed from minimap2 mappings, and consecutively sequenced reads with adjacent alignment within a genomic distance of –10 nt to 1,000 nt were identified. Samples sequenced before or during May 2018 had very low levels of over-splitting (between 0.01% and 0.05% of reads) compared with those sequenced in September 2018 onwards (between 0.8% and 1.5% of reads).

#### Analysis of the potential for internal priming in nanopore DRS data

To determine whether internal priming caused by the RT step can occur in nanopore data, the location of oligo(A) hexamers within Arabidopsis coding sequence (CDS) regions (Araport11) was determined and reads that terminated within a 20 nt window of each hexamer were counted. Of the 10,116 CDS oligo(A) runs, 160 (1.58%) had at least one supporting read in one Col-0 nanopore dataset. Of these, 137 were supported by only one replicate, and only four were supported by all four biological replicates. In total, 66 (41%) occurred in Araport11-annotated terminal exons, suggesting that they may be genuine sites of 3’ end formation.

#### Poly (A) length estimation using nanopore DRS data

Poly (A) tail length estimations were produced using nanopolish version 0.11.0^18^ and added as tags to BAM files using pysam version 0.15.2^62^. Per gene length distributions were then produced using Araport11 annotation, and genes with significant changes in length distribution in the *vir-1* mutant compared with the VIR-complemented line were identified using a Kolmogorov–Smirnov test*. p*-values were adjusted for multiple testing using Benjamini-Hochberg correction.

#### 3′ end analyses of nanopore and Helicos reads

Helicos data were prepared as previously described^20, 63^. Positions with three or more supporting reads were considered to be peaks of nanopore or Helicos 3′ ends. The distance between each nanopore peak and the nearest Helicos peak was then determined. In all, 37% of nanopore peaks occurred at the same position as a Helicos peak, and the standard deviation in distance was 12.5 nt. To determine the percentage of nanopore DRS 3′ ends mapping within annotated genic features, transcripts were first flattened into a single record per gene. Exonic annotation was given priority over intronic or intergenic annotation and CDS annotation was given priority over UTR annotation. Reads were assigned to genes if they overlapped them by >20% of their aligned length, and the annotated feature type of the 3′ end position was determined. Counts were generated both for all reads and for unique positions per gene.

#### Isoform collapsing of nanopore DRS data

Error-corrected full-length alignments were collapsed into clusters of reads with identical sets of introns. These clusters were then subdivided by 3′ end location by using a Gaussian kernel with sigma of 100 to find local minima between read ends, which were used as cut points to separate clusters. The read with the longest aligned length in each cluster was used as the representative in the figure.

#### Splicing analysis of nanopore DRS and Illumina RNAseq data

Splice junction locations, their flanking sequences and the read counts supporting them were extracted from Illumina RNA sequencing, nanopore DRS and nanopore error-corrected DRS reads using pysam version 0.14^62^, as well as from Araport11^64^ and AtRTD2^25^ reference annotations. Splice junctions at the same position but on the opposite strand were counted independently. Junctions were classified by their most likely snRNP machinery using Biopython version 1.71^65^, with position weight matrices as previously calculated^28^. Position weight matrices were scored against the sequence –3 nt to +10 nt of the donor site and –14 nt to +3 nt of the acceptor site. Position weight matrix scores greater than zero indicate a match to the motif, while scores of around zero, or negative scores, indicate background frequencies or deviation from the motif. Positive scores were normalized into the range 50–100 as done previously^28^. Junctions with U12 donor scores of >75 and U12 acceptor scores of >65 were classified as U12 junctions, while junctions with U2 donor and acceptor scores of >60 were classified as U2, as done previously^25^. Junctions were further categorized as canonical or non-canonical based on the presence or absence, respectively, of GT/AG intron border sequences. For isoform analysis, linked splices from the same read were extracted from full-length nanopore error-corrected reads and counted to create unique sets of splice junctions. Intronless reads were not counted. UpSet plots were generated in Python 3.6 using the package upsetplot^66^.

#### Validation of novel splice sites

To validate novel splice junctions detected in nanopore DRS, five splice sites out of the 20 most highly expressed splice sites were selected for further validation; three of the five selected splice sites were successfully amplified in RT-PCR followed by Sanger sequencing (described above).

#### 5′ adapter detection analyses using nanopore DRS data

To produce positive and negative examples of 5′ adapter-containing sequences, 5′ soft-clipped regions were extracted from aligned reads for the Col-0 replicate 1 datasets (with and without adapter ligation) using pysam^62^. These soft-clipped sequences were then searched for the presence of the GeneRacer adapter sequence using BLASTN version 2.7.1^67^. Two rules were initially applied to filter BLASTN results: a match of 10 nt or more to the 44 nt adapter, and an E value of <100. Reads from the adapter-containing dataset that failed one or both criteria were used as negative training examples. A final rule requiring the match to the adapter sequence to occur directly adjacent to the aligned read was also applied. Reads from the adapter-containing dataset that passed all three rules were used as the positive training set. When comparing the ratio of positive to negative examples between datasets containing the adapter and those generated from the same tissue but without the adapter, we found that these three rules gave a signal-to-noise ratio of >5,000 (Supplementary table 2).

In all, 72,083 positive and 123,739 negative training examples from Col-0 tissue replicate 1 were collected to train the neural network. We then estimated the amount of raw signal from the 5′ end of the squiggle that was required on average to capture the 5′ adapter. To do this, we used nanopolish eventalign version 0.11.0^68^ to identify the interval in the raw read corresponding to the mRNA alignment to the reference in the positive examples of 5′ adapter-containing sequences. Since the adapter can be identified immediately adjacent to the alignment in sequence space for these reads, the signal after the event alignment should correspond to the signal originating from the adapter. The median length of these signals was 1,441 points, and 96% of the signals were <3,000 points. Therefore, we used a window size of 3,000 to make predictions.

The model was trained in Python 3.6 using Keras version 2.2.4 with the Tensorflow version 1.10.0 backend^69, 70^. A ResNet-style architecture was used^71^, composed of eight residual blocks containing two convolutional layers of kernel size 5 and a shortcut convolution with kernel size 1. Down-sampling using maximum pooling layers with a stride of 2 was used between each residual block. A penultimate densely connected layer of size 16 was used, with training dropout of 0.5. Input signals were standardized by median absolute deviation scaling across the whole read before the final 3,000 points were taken, and the negative samples were augmented by addition of random internal signals from reads and pure Gaussian, multi-Gaussian and perlin noise signals^72^. The whole dataset was also augmented on the fly during training by the addition of Gaussian noise with a standard deviation of 0.1. Models were trained for a maximum of 100 epochs (batch size of 128, 100 batches per epoch, positive and negative examples sampled in a 1:1 ratio) using the RMSprop optimizer with an initial learning rate of 0.001, which was reduced by a factor of 10 after three epochs with no reduction in validation loss. Early stopping was used after five epochs with no reduction in validation loss. We found that a number of negative training examples from the ends of reads, but not from internal positions, were likely to be incorrectly labelled by the BLASTN method, because the model predicted them to contain adapters. We therefore filtered these to clean the training data, before repeating the training process. Model performance was evaluated using five-fold cross validation and by testing on independently generated datasets from Col-0 replicate 2, produced with and without the adapter ligation protocol (Supplementary figure 3B, C)^69, 70^.

To evaluate the reduction in 3′ bias of adapter-ligated datasets, we used Araport11 exon annotations to produce per base coverage for each gene in the Col-0 replicate 2 dataset. Coverage was generated separately for reads predicted to contain adapters and for those predicted not to contain adapters. Leading zeros at the extreme 5′ and 3′ ends of genes were assumed to be caused by mis-annotation of UTRs and so were trimmed. The quartile coefficient of variation (interquartile range / median) was then used as a robust measure of variation in coverage across each gene. To validate the 5′ ends of adapter-ligated reads with orthogonal data, full-length cDNA clone sequences were downloaded from RIKEN RAFL (Arabidopsis full-length cDNA clones)^23^. These were mapped with minimap2^13^ in spliced mode. The distance from each nanopore alignment 5′ end to the nearest RIKEN RAFL alignment^23^. 5′ end was calculated using bedtools^58^. The amount of 5′ end sequence that is rescued when 5′ adapters are used was estimated by identifying the largest peak in 5′ end locations per gene in the absence of adapter, and then measuring how this peak shifted using reads predicted to contain adapters.

#### Differential error site analysis using nanopore DRS data

To detect sites of Virilizer-dependent m^6^A RNA modifications, we developed scripts to test changes in per base error profiles of aligned reads. Pileup columns for each position with coverage of >10 reads were generated using pysam^62^ and reads in each column were categorized as either A, C, G, T or indel. The relative proportions of each category were counted. Counts from replicates of the same experimental condition were aggregated and a 2×5 contingency table was produced for each base comparing *vir-1* and VIR-complemented lines. A G-test was performed to identify bases with significantly altered error profiles. For bases with a *p-*value of <0.05, G-tests for homogeneity between replicates of the same condition were then performed. Bases where the sum of the G statistic for homogeneity tests was greater than the G statistic for the *vir-1* and VIR-complemented line comparison were filtered. Multiple testing correction was carried out using the Benjamini-Hochberg method, and an FDR threshold of 0.05 was used. The log2 fold change in mismatch to match ratio (compared with the reference base) between the *vir-1* and VIR-complemented lines was calculated using Haldane correction for zero counts. Bases that had a log fold change of >1 were considered to have a reduced error rate in the *vir-1* mutant.

To identify motifs enriched at sites with a reduced error rate, reduced error rate sites were increased in size by 5 nt in each direction and overlapping sites were merged using bedtools version 2.27.1^58^. Sequences corresponding to these sites were extracted from the TAIR10 reference and over-represented motifs were detected in the sequences using MEME version 5.0.2^73^, run in zero or one occurrence mode with a motif size range of 5–7 and a minimum requirement of 100 sequence matches. The presence of these motifs at error sites was then detected using FIMO version 5.0.2^74^. A relaxed FDR threshold of 0.1 was used with FIMO to capture more degenerate motifs matching the m^6^A consensus.

#### Differential gene expression analysis using nanopore DRS data

Gene level counts were produced for each nanopore DRS replicate using featureCounts version 1.6.3^75^ in long-read mode with strand-specific counting. Differential expression analysis between the *vir-1* and VIR-complemented lines was then performed in R version 3.5 using edgeR version 3.24.3^55^.

#### Identification of alternative 3′ end positions and chimeric RNA using nanopore DRS data

Genes with differential 3′ end usage were identified by producing 3′ profiles of reads which overlapped with each annotated gene locus by >20%. These profiles were then compared between the *vir-1* and VIR-complemented lines using a Kolmogorov–Smirnov test to identify changes. Multiple testing correction was performed using the Benjamini-Hochberg method. To approximately identify the direction and distance of the change, the normalized single base level histograms of the VIR-complemented line profile was subtracted from that of the mutant profile, and the minimum and maximum points in the difference profile were identified. These represent the sites of most reduced and increased relative usage, respectively. Results were filtered for an FDR of <0.05 and absolute change of site >13 nt (the measured error range of nanopore DRS 3′ end alignment). The presence of m^6^A modifications at genes with differential 3′ end usage was assessed using bedtools intersect^58^, and significant enrichment of m^6^A at these genes was tested using a hypergeometric test (using all expressed genes as the background population).

To identify genes with a significant increase in chimeras in the *vir-1* mutant, we used Araport11 annotation^64^ to identify reads that overlapped with multiple adjacent gene loci (i.e. chimeric reads) and those originating from a single locus (i.e. non-chimeric reads). To reduce false positives caused by reads mapping across tandem duplicated loci, reads mapping to genes annotated in PTGBase^76^ were filtered out. Chimeric reads were considered to originate from the most upstream gene with which they overlapped. We pooled reads from replicates for each experimental condition and used 50 bootstrapped samples (75% of the total data without replacement) to estimate the ratio of chimeric to non-chimeric reads at each gene in each condition. Haldane correction for zero counts was applied. The distributions of chimeric to non-chimeric ratios in the *vir-1* and VIR-complemented lines were tested using a Kolmogorov–Smirnov test to detect loci with altered chimera production. All possible pairwise combinations of VIR-complemented and *vir-1* bootstraps were then compared to produce a distribution of estimates of change in the chimeric to non-chimeric ratio in the *vir-1* mutant. Loci that had more than one chimeric read in *vir-1*, demonstrated at least a two-fold increase in the chimeric read ratio in >50% of bootstrap comparisons and were significantly changed at a FDR of <0.05 were considered to be sites of increased chimeric RNA formation in the *vir-1* mutant.

### miCLIP

#### Preparation of miCLIP libraries

Total RNA for miCLIP was isolated from 7.5 mg of 14-day old Arabidopsis Col-0 seedlings as previously described^77^. mRNA was isolated from ∼1 mg total RNA using oligo(dT) and streptavidin paramagnetic beads (PolyATtract mRNA Isolation Systems, Promega) according to the manufacturer’s instructions. miCLIP was carried out using 15 µg mRNA as previously described^78^ using an antibody against N6-methyladenosine (#202 003 Synaptic Systems), with minor modifications. No-antibody controls were processed throughout the experiment. RNA-antibody complexes were separated by 4–12% Bis-Tris gel electrophoresis at 180 V for 50 min and transferred to nitrocellulose membranes (Protran BA85 0.45µm, GE Healthcare) at 30 V for 60 min. Membranes were then exposed to Medical X-Ray Film Blue (Agfa) at −80°C overnight. Reverse transcription was carried out using barcoded RT primers: RT41, RT48, RT49 and RT50 (Integrated DNA Technologies; Supplementary table 5). After reverse transcription, one cDNA fraction corresponding to 70–200 nt was gel purified after 6% TBE-urea gel electrophoresis (Thermo Fisher Scientific). After the final PCR step, all libraries were pooled together, purified using Agencourt Ampure XP magnetic beads (Beckman Coulter) and eluted in nuclease-free water. Paired-end sequencing with a read length of 100 bp was carried out on an Illumina MiSeq v2 at Edinburgh Genomics, University of Edinburgh. Input sample libraries were prepared using NEBNext Ultra Directional RNA Library Prep Kit for Illumina (New England Biolabs) and sequenced on an Illumina HiSeq2000 at the Tayside Centre for Genomics Analysis, University of Dundee, with a pair-end read length of 75 bp.

#### Processing of miCLIP sequencing data

miCLIP data were assessed for quality using FastQC version 0.11.8^79^ and MultiQC version 1.7^80^. Only the forward read was used for analysis because the miCLIP site is located at the 5′ position of the forward read. 3′ adapter and poly(A) sequences were trimmed using cutadapt version 1.18^81^ and unique molecular identifiers were extracted from the 5′ end of the reads using UMI-tools version 0.5.5^82^. Immunoprecipitation and no-antibody controls were demultiplexed and multiplexing barcodes were trimmed using seqkit version 0.10.0^59^. Reads were mapped to the TAIR10 reference genome with Araport11 reference annotation^64^ using STAR version 2.6.1^52^, a maximum multimapping rate of 5, a minimum splice junction overhang of 8 nt (3 nt for junctions in the Araport11 reference), a maximum of five mismatches per read, and intron length boundaries of 60– 10,000 nt. SAM and BAM file manipulations were performed using samtools version 1.9^56^. Removal of PCR duplicates was then performed using UMI-tools in a directional model^82^. miCLIP 5′ coverage and matched input 5′ coverage tracks were generated using bedtools version 2.27.1^58^ and these were used to call miCLIP peaks at single nucleotide resolution with Piranha version 1.2.1^83^ and relaxed *p*-value thresholds of 0.5. Reproducible peaks across pairwise combinations of the three replicates were identified by irreproducible discovery rate (IDR) analysis using Python package idr version 2.0.3 with an IDR threshold of 0.05^84^. The final set of peaks was identified by pooling the three replicates, re-analysing using Piranha, ranking the peaks by FDR and selecting the top N peaks, where N is the smallest number of reproducible peaks discovered by pairwise comparisons of the three replicates. This yielded 141,198 unique nucleotide-level miCLIP peaks.

### m^6^A liquid chromatography–mass spectroscopy analysis

m^6^A content analysis using liquid chromatography–mass spectroscopy (LC-MS) was performed as previously described^85^. Chromatography was carried out by the FingerPrints Proteomics facility, University of Dundee.

## Code availability

All scripts, pipelines and notebooks used for this study are available on GitHub at https://github.com/bartongroup/Simpson_Barton_Nanopore_1

## Data availability

Sequencing datasets described in this study have been deposited at the European Nucleotide Archive understudy accession number PRJEB32782.

## Acknowledgements

This work was funded by BBSRC grants BB/J00247X/1, BB/M004155/1, BB/M010066/1, the University of Dundee Global Challenges Research Fund and the European Union’s Horizon 2020 research and innovation programme under Marie Skłodowska-Curie grant agreement No. 799300. The m^6^A LC-MS analysis was carried out by Abdel Atrih of the FingerPrints Proteomics Facility, University of Dundee, which is supported by a Wellcome Trust Technology Platform award (097945/B/11/Z). We thank Kasper Rasmussen and Csaba Hornyik for their comments on the manuscript.

## Author contributions

GGS, NJS and AVS conceived the study; KK, AVS and KM designed and performed the experiments; MTP and NJS undertook data analysis; PDG undertook the clock experiments, supervised by AH; GGS and GJB supervised the study; GGS, MTP and AVS wrote the paper. All authors read and commented on the text.

---

**Supplementary figure 1.**
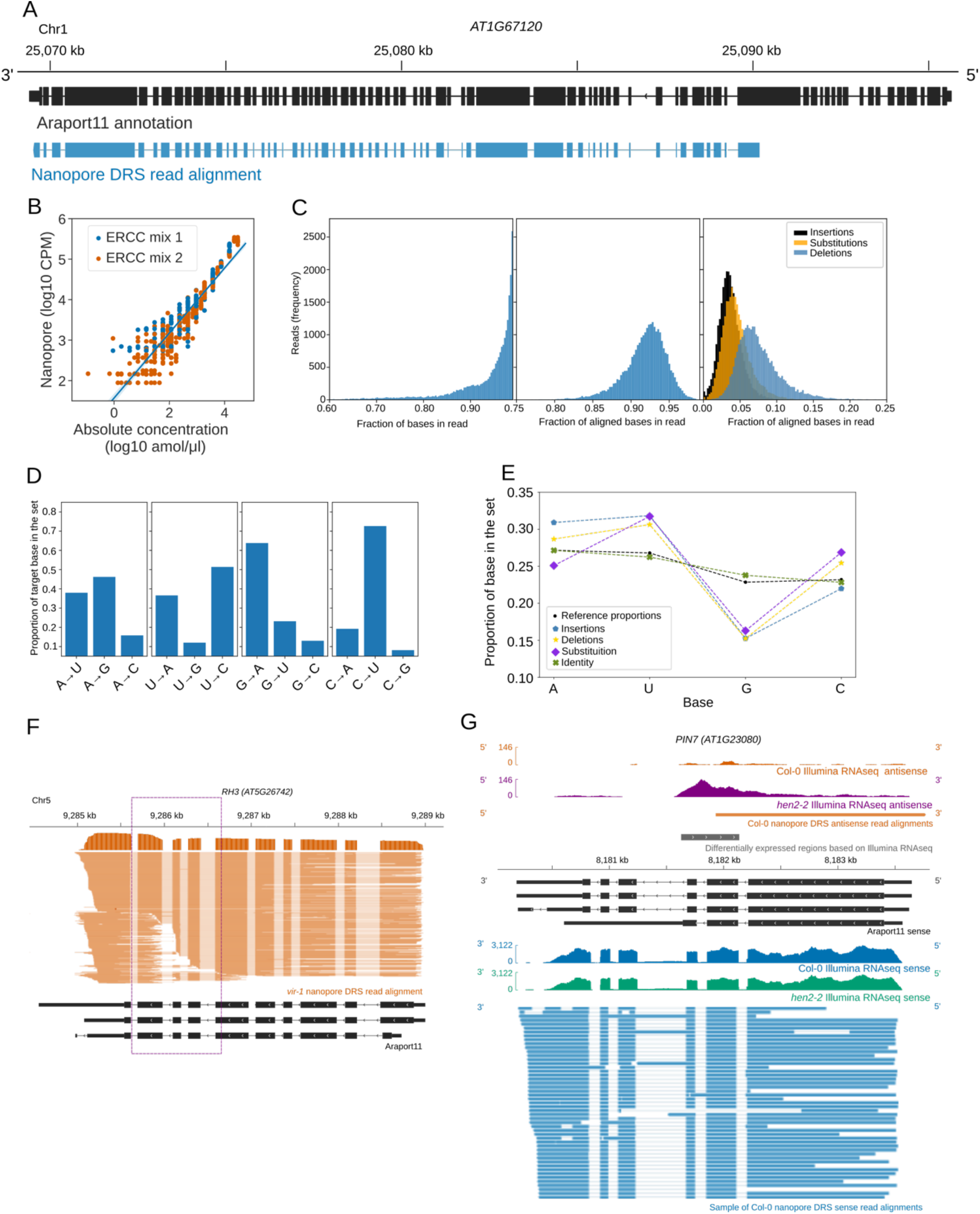
Properties of nanopore DRS sequencing data. (**A**) Nanopore DRS identified a 12.8 kb transcript generated from the *AT1G67120* gene that includes 58 exons. Black, RNA isoform present in the Araport11 annotation; blue, an RNA isoform identified using nanopore DRS. (**B**) Synthetic ERCC RNA Spike-In mixes are detected in a quantitative manner. Absolute concentrations of spike-ins are plotted against counts per million (CPM) reads in log10 scale. Blue, ERCC RNA Spike-In mix 1; orange, ERCC RNA Spike-In mix 2. (**C**) Overview of the sequencing and alignment characteristics of nanopore DRS data for ERCC RNA Spike-Ins. Left, distribution of the length fraction of each sequenced read that aligns to the ERCC RNA Spike-In reference; centre, distribution of fraction of identity that matches between the sequence of the read and the ERCC RNA Spike-In reference for the aligned portion of each read; right, distributions of the occurrence of insertions (black), substitutions (orange) and deletions (blue) as a proportion of the number of aligned bases in each read. (**D**) Substitution preference for each nucleotide (left to right: adenine [A], uracil [U], guanine [G], cytosine [C]). When substituted, G is replaced with A in more than 63% of its substitutions, while C is replaced by U in 73%. Conversely, U is rarely replaced with G (12%) and A is rarely substituted with C (16%). (**E**) Nucleotide representation within the ERCC RNA Spike-In reference sequences (black dots) compared with nucleotide representation within four categories from the nanopore DRS reads. Identity matches between the sequence of the read and the ERCC RNA Spike-In reference (green crosses), insertions (blue pentagons), deletions (yellow stars) and substitutions (purple diamonds).G is under-represented and U is over-represented for all three categories of error (insertion, deletion and substitution) relative to the reference nucleotide distribution. C is over-represented in deletions and substitutions. A is over-represented in insertions and deletions and under-represented in substitutions. (**F**) Signals originating from the *RH3* transcripts are susceptible to systematic over-splitting around exons 7–9 (highlighted using a purple dashed box), resulting in reads with apparently novel 5′ or 3′ positions. This appears only to occur at high frequency in datasets collected after May 2018 (Supplementary table 1) and may result from an update to the MinKNOW software. (**G**) *PIN7* long non-coding antisense RNAs detected using nanopore DRS. Blue, Col-0 sense Illumina RNAseq coverage and nanopore sense read alignments; orange, Col-0 antisense Illumina RNAseq coverage and nanopore antisense read alignments; green, *hen2–2* mutant sense Illumina RNAseq coverage; purple, *hen2–2* mutant antisense Illumina RNAseq coverage; black, sense RNA isoforms found in Araport11 annotation; grey, antisense differentially expressed regions detected with DERfinder. **[Linked to Figure 1]**

**Supplementary figure 2.**
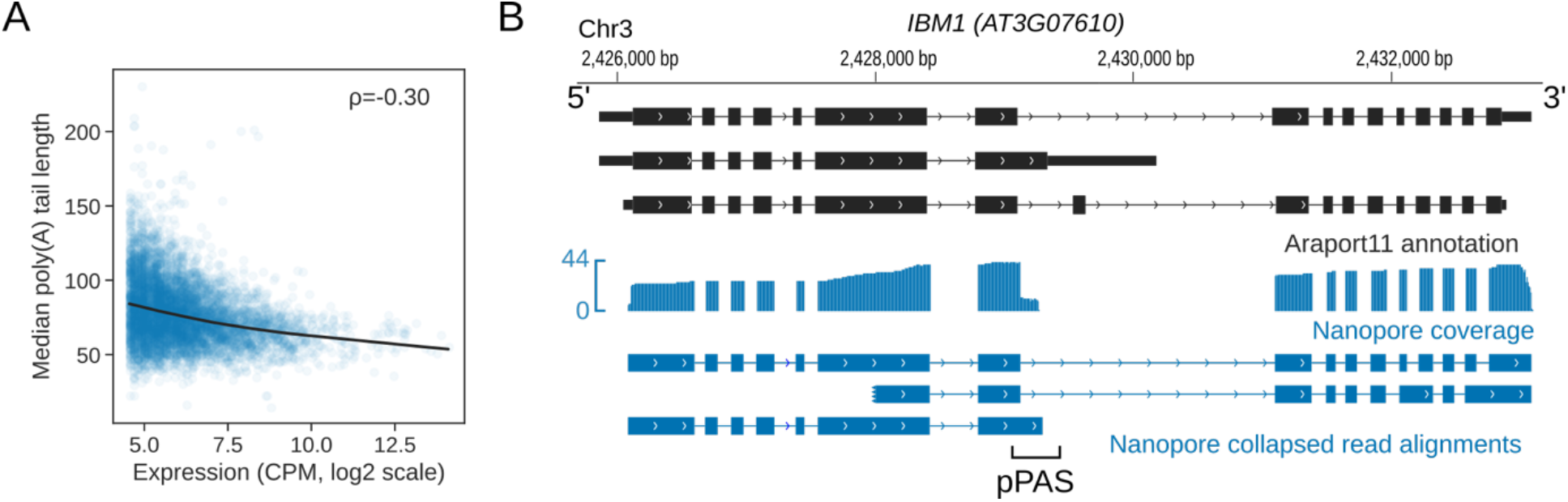
3′ end processing is revealed by nanopore DRS. (**A**) The RNA poly(A) tail length negatively correlates with the gene expression level. Expression in log2 scale of counts per million (CPM) obtained from nanopore DRS data is plotted against the median poly(A) tail length. ρ, Spearman’s correlation coefficient; black line, locally weighted scatterplot smoothing (LOWESS) regression fit. (**B**) Nanopore DRS identified 3′ polyadenylation sites in RNAs transcribed from the *IBM1 (AT3G07610)* gene. The blue track shows the coverage of nanopore DRS reads. Black, isoforms found in Araport11 annotation; blue, isoforms those detected by nanopore DRS. pPAS, proximal polyadenylation site. **[Linked to Figure 2].**

**Supplementary figure 3.**
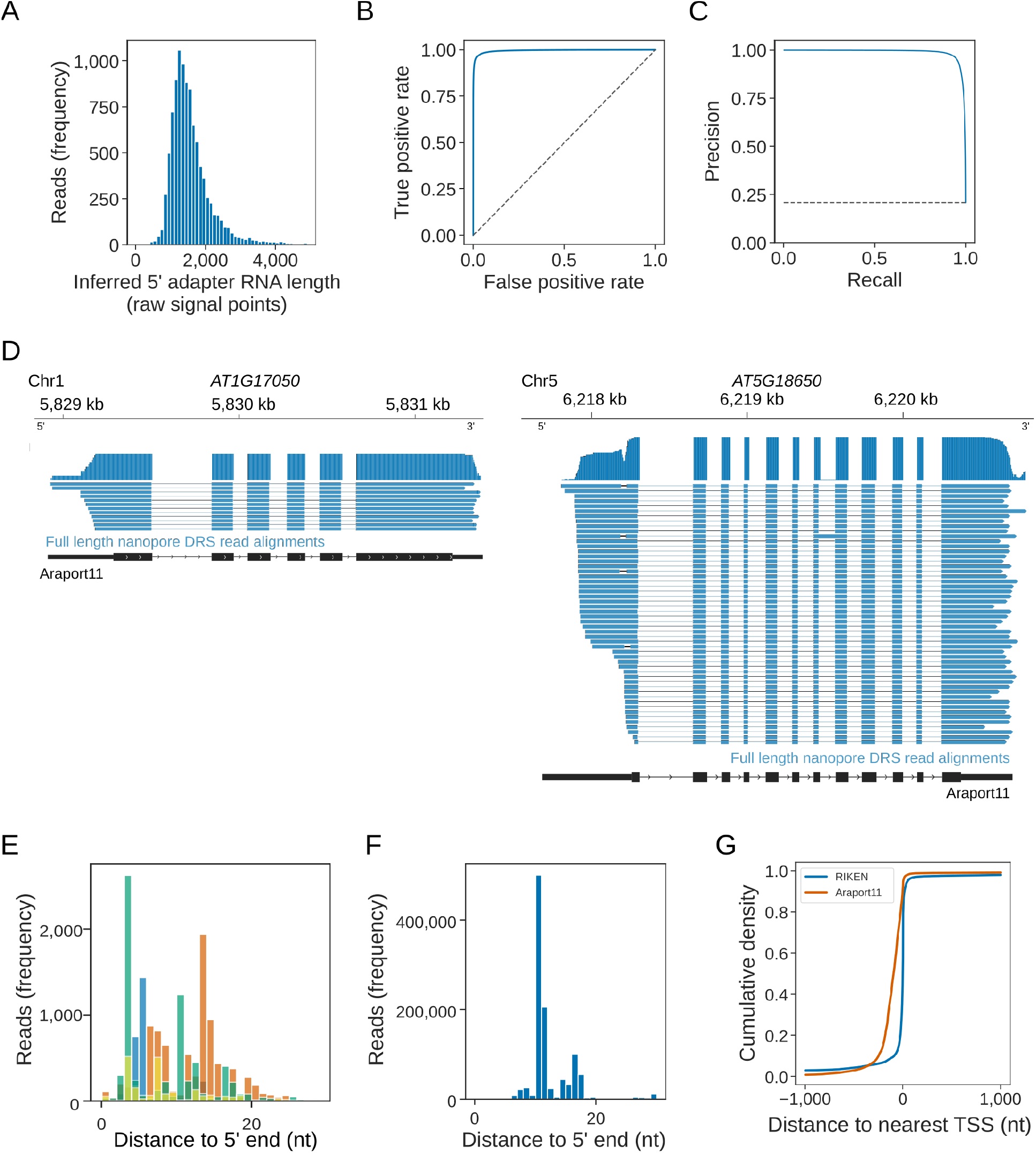
Nanopore DRS with cap-dependent ligation of 5′ adapter RNA. (**A**) Histogram showing the distribution of 5′ adapter RNA length in the nanopore raw current signal, as inferred from alignment of the mRNA sequence to the signal using nanopolish eventalign. The median signal length was 1,441 points and 96% of adapter signals were 3,000 points or less. (**B**) Out-of-bag receiver operator characteristic curve showing the performance of the trained convolutional neural network at detecting 5′ adapter RNA using 3,000 points of signal. The curve was generated using five-fold cross validation. (**C**) Out-of-bag precision recall curve showing the performance of trained neural network, generated using five-fold cross validation. (**D**) Alternative transcription start sites were identified using nanopore DRS with cap-dependent ligation of a 5′ end adapter at the *AT1G17050* and *AT5G18650* genes. The blue track shows the coverage of nanopore DRS reads. Black, isoforms found in the Araport11 annotation; blue, isoforms detected by nanopore DRS with cap-dependent ligation of 5′ adapter RNA. (**E**) Reads mapping to ERCC RNA Spike-Ins lack approximately 11 nt of sequence at the 5′ end. Histogram showing the distance to the 5′ end for ERCC RNA Spike-In reads (each spike-in is shown in a different colour; only those with >1,000 supporting reads are shown). (**F**) Reads mapping to *in vitro* transcribed mGFP lack approximately 11 nt of sequence at the 5′ end. Histogram showing the distance to the 5′ end for *in vitro* transcribed mGFP. (**G**) Araport11 annotation overestimates the length of 5′ UTRs. The cumulative distribution function shows the distance to the nearest TSS identified from full-length transcripts cloned as part of the RIKEN RAFL project (blue) and Araport11 annotation (orange). **[Linked to Figure 3].**

**Supplementary figure 4.**
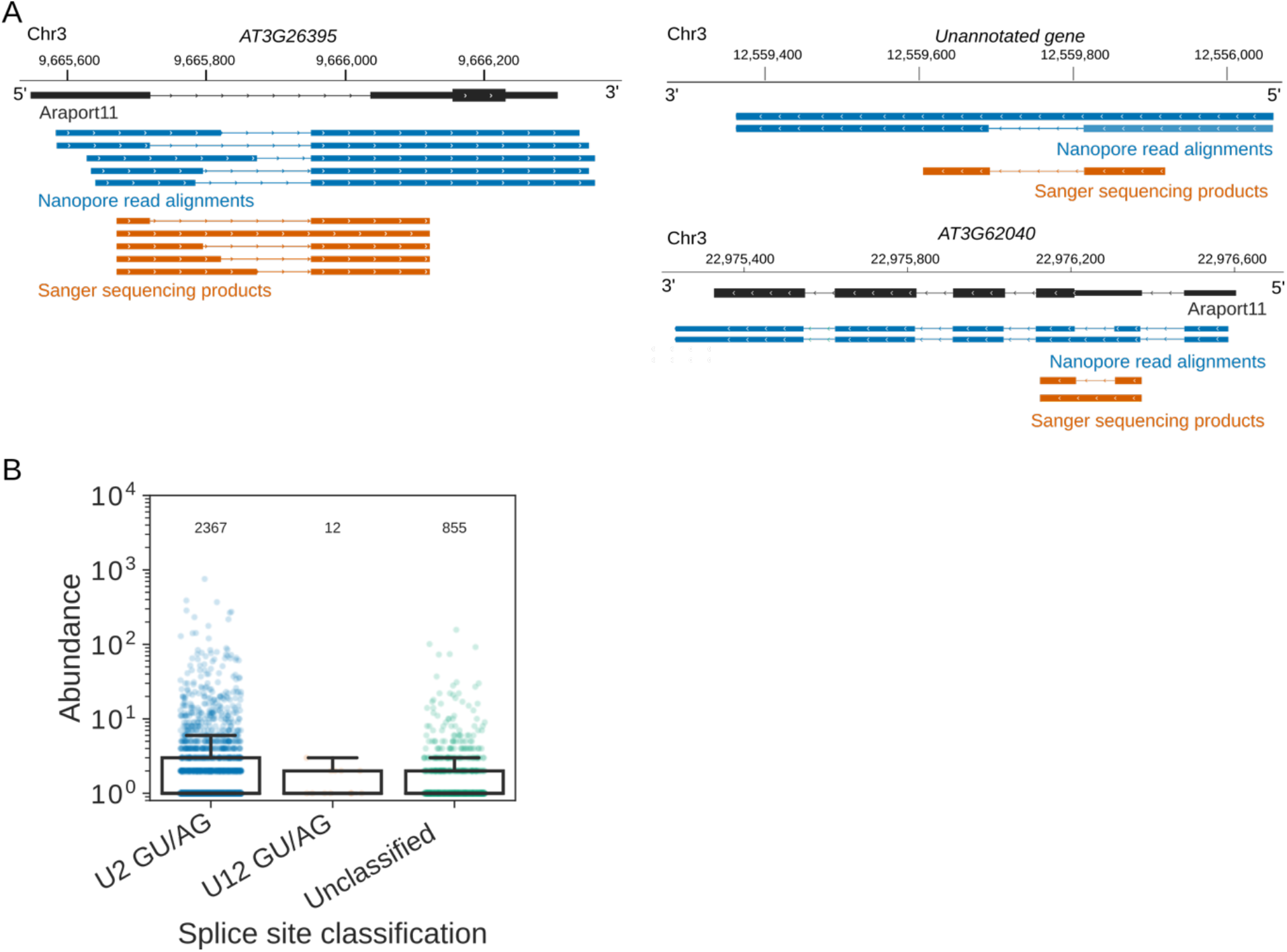
Patterns of splicing revealed using nanopore DRS. (**A**) Nanopore DRS can be validated using RT-PCR. Five of the top 20 most highly expressed RNAs with novel splice sites were selected; of these, three were validated by RT-PCR followed by Sanger sequencing of the DNA products. Black, RNA isoforms present in the Araport11 annotation; blue, RNA isoforms found using nanopore DRS; orange, Sanger sequencing products obtained using RT-PCR. (**B**) Splice junction classification of unannotated GU/AG splice sites found in error-corrected nanopore DRS data that also have Illumina support. Counts are plotted in log10 scale and the exact number of splice junctions in each set is indicated. **[Linked to Figure 4].**

**Supplementary figure 5.**
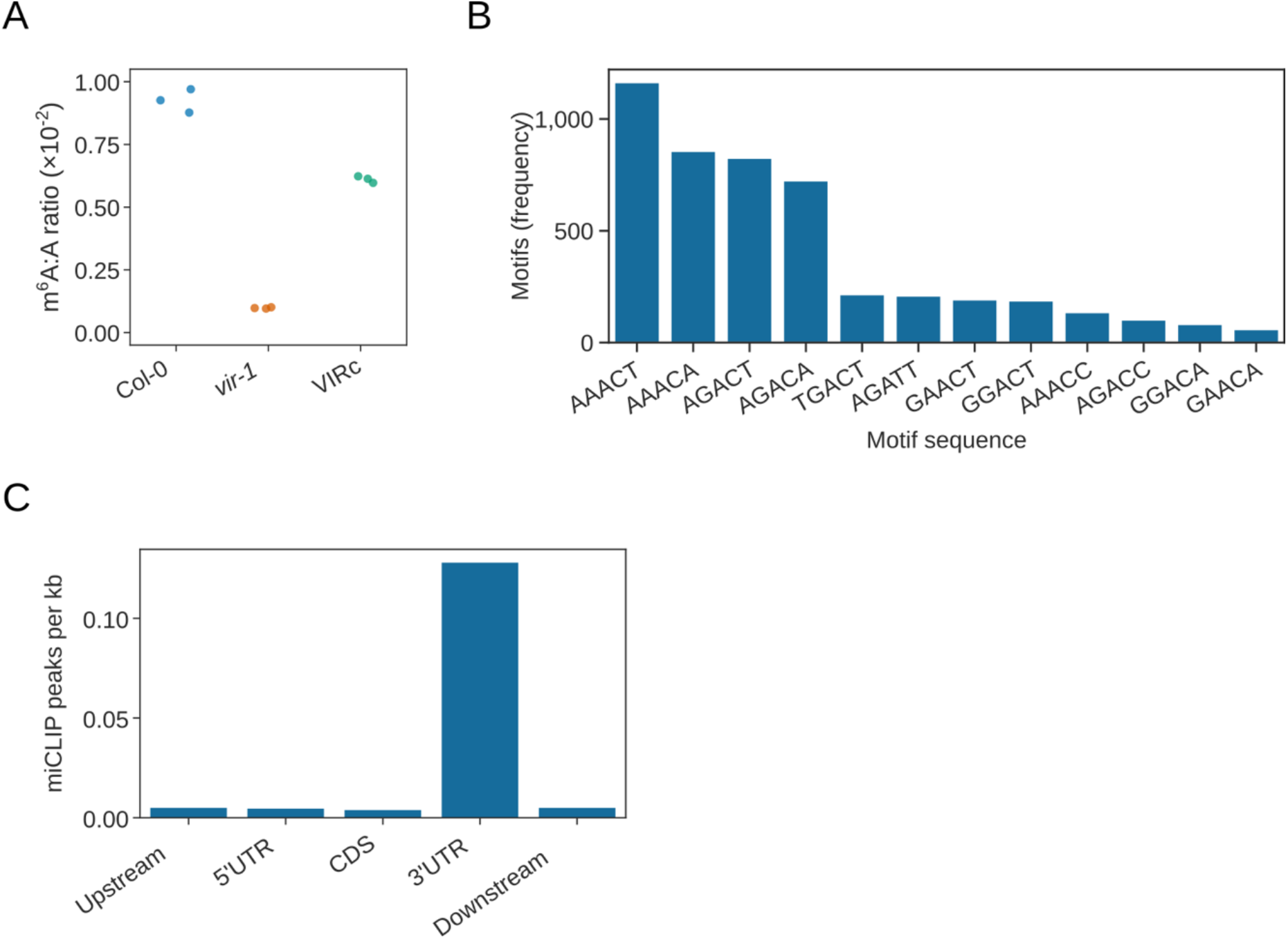
Identification of VIR-dependent m^6^A transcriptome-wide. (**A**) *vir-1* shows reduced levels of m^6^A compared with Col-0 and restored m^6^A levels in the VIR-complemented line (VIRc). The ratio of m^6^A/A obtained using LC-MS analysis is shown Col-0 (blue), *vir-1* (orange) and VIRc (green). (**B**) Frequency of m^6^A motifs detected at *vir-1* reduced error sites, as detected by FIMO using the motif detected *de novo* by MEME and an FDR threshold of 0.1. (**C**) Bar plot showing the number of miCLIP peaks per kb of different genic feature types in the Araport11 reference. Upstream and downstream regions were defined as 200 nt regions before and after the annotated transcription termination sites, respectively. **[Linked to Figure 5].**

**Supplementary figure 6.**
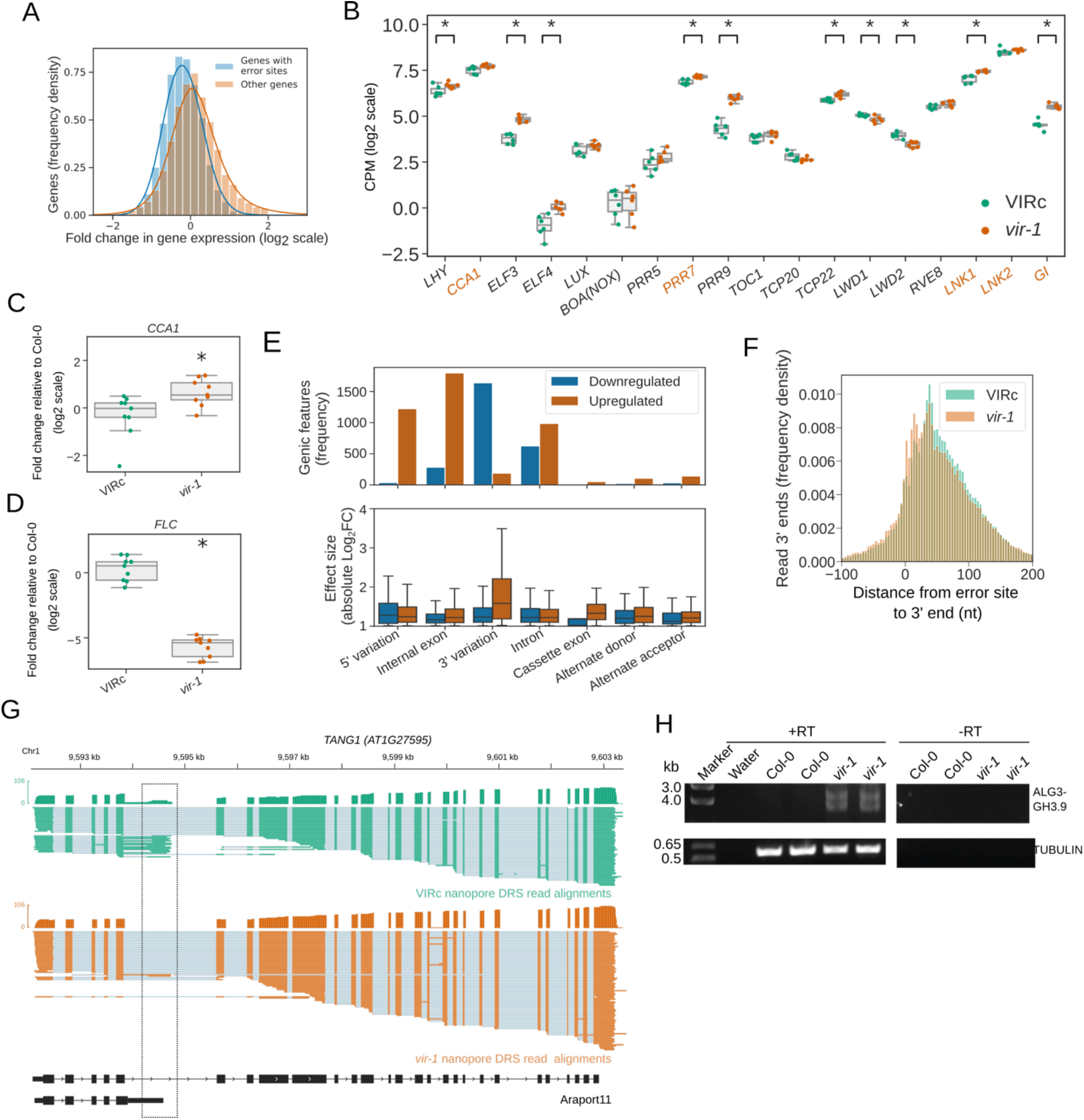
Changes in the gene expression of circadian clock components and in RNA 3′ end formation in the *vir-1* mutant. (**A**) Histogram showing log2 fold changes in gene expression based on Illumina RNAseq data for the *vir-1* mutant and VIR-complemented line. Blue, genes with differential error rate sites (n=5,169 genes); orange, genes without differential error rate sites (n=14,612 genes). (**B**) Expression of core circadian clock components is perturbed in the *vir-1* mutant. Boxplots showing normalized gene expression measured using Illumina RNAseq in log2 counts per million (CPM): green, the VIR-complemented line (VIRc); orange, the *vir-1* mutant. Each scatter point represents a single biological replicate. Asterisks denote significant expression changes (using an FDR threshold of 0.05). Orange labelled genes have 3′ UTR m^6^A detectable by nanopore DRS and miCLIP. (**C**) Expression of *CCA1*, encoding a regulator of the circadian rhythm in Arabidopsis, is increased in the *vir-1* mutant. Boxplot showing the gene expression change from Col-0 (measured by RT-qPCR) for VIRc (green) and *vir-1* (orange). Three technical replicates of three biological replicates were conducted. Each scatter point represents the comparison of a technical replicate of treatment (VIRc or *vir-1*) against control (Col-0). The expression change in *vir-1* is significant (*p*=0.02). (**D**) Expression of the *Flowering Locus C* (*FLC*) gene is decreased in the *vir-1* mutant. Boxplot showing gene expression change from Col-0 (measured by RT-qPCR) for VIRc (green) and *vir-1* (orange). Three technical replicates of three biological replicates were conducted. Each scatter point represents the comparison of a technical replicate of treatment (VIRc or *vir-1*) against control (Col-0). Expression change in *vir-1* is significant (*p*=2.4×10^-14^). (**E**) Splicing is moderately disrupted in the *vir-1* mutant. Results of differential exon usage analysis with DEXseq are shown for contiguous regions (“exon chunks”), which occur in the same sets of transcripts in the Araport11 reference. Regions were classified as a 5′ or 3′ variation if they were bounded by the TSS of one or more transcripts. Orange, features with increased usage; blue, features with reduced usage. Boxplots show the distribution in absolute log2 fold change for each feature set. (**F**) A shift to the use of more proximal polyadenylation sites is observed in m^6^A containing transcripts in the *vir-1* mutant. Histogram showing distance from the error site (n=17,491 error sites) to upstream and downstream 3′ ends in the *vir-1* mutant (orange) and VIRc (green). (**G**) *vir-1* mutants exhibit increased readthrough of an intronic proximal poly(A) site in intron 5 of the Symplekin domain encoding gene *TANG1*. green, nanopore DRS reads from VIRc; orange, nanopore DRS reads from the *vir-1* mutant; black, Araport11 annotation. The dashed black box highlights the site of proximal polyadenylation. (**H**) *ALG3-GH3.9* chimeric RNAs are generated in the *vir-1* mutant. RT-PCR gel showing formation of chimeric RNAs in the *vir-1* mutant compared with Col-0. **[Linked to Figure 6].**

**Supplementary table 1.** Properties of the nanopore DRS sequencing data. Dataset statistics for all nanopore DRS sequencing runs conducted. Datasets are sorted by the date of the sequencing run. All data was collected using a MinION with R9.4 flow cell and SQK-RNA001 library kit. Increases in mapping and over-splitting rate that occur in samples collected after September 2018 are therefore likely to have resulted from changes in the MinKNOW software. **[Linked to Figure 1 through 6].**

| Sequencing date | Genotype | Bio replicates | 5' Adapter ligation | Total reads basecalled | Reads mapped to ERCC spike-ins | Reads mapped to TAIR10 | Longest read mapped to ERCC spike-ins | Median length of reads mapped to ERCC spike-ins | Longest read mapped to TAIR10 | Median length of reads mapped to TAIR10 | Percentage mapped to TAIR10 | Estimated over-splitting rate (%) |
| --- | --- | --- | --- | --- | --- | --- | --- | --- | --- | --- | --- | --- |
| 01/02/2018 | Col-0 | 1 | - | 1,647,484 | 6756 | 1,003,137 | 1,905 | 481 | 12,607 | 865 | 60.89 | 0.047 |
| 27/02/2018 | vir-1 | 1 | - | 1,431,457 | 1074 | 1,005,664 | 2,304 | 492 | 12,446 | 878 | 70.25 | 0.045 |
| 01/03/2018 | VIRc | 1 | - | 1,393,351 | 713 | 1,101,368 | 1,108 | 493 | 11,994 | 873 | 79.04 | 0.111 |
| 05/04/2018 | Col-0 | 2 | - | 966,529 | 556 | 743,684 | 1,099 | 492 | 11,852 | 878 | 76.94 | 0.035 |
| 11/04/2018 | Col-0 | 1 | + | 1,365,809 | 7005 | 345,799 | 1,928 | 486 | 5,663 | 737 | 25.32 | 0.047 |
| 13/04/2018 | Col-0 | 2 | - | 1,242,616 | 701 | 930,629 | 1,094 | 488 | 11,434 | 858 | 74.89 | 0.040 |
| 16/04/2018 | Col-0 | 3 | - | 1,079,578 | 907 | 757,028 | 1,772 | 486 | 12,744 | 840 | 70.12 | 0.019 |
| 18/04/2018 | Col-0 | 4 | - | 1,007,278 | 525 | 765,322 | 1,292 | 495 | 11,542 | 840 | 75.98 | 0.019 |
| 08/05/2018 | Col-0 | 2 | + | 2,043,751 | 11324 | 557,949 | 2,308 | 507 | 5,596 | 732 | 27.30 | 0.069 |
| 07/09/2018 | VIRc | 2 | - | 1,699,123 | 1824 | 1,538,040 | 2,236 | 497 | 12,015 | 871 | 90.52 | 0.809 |
| 12/09/2018 | vir-1 | 2 | - | 858,747 | 1094 | 812,517 | 1,880 | 491 | 15,040 | 846 | 94.62 | 0.959 |
| 25/09/2018 | VIRc | 4 | - | 1,435,808 | 2710 | 1,318,857 | 1,947 | 493 | 10,836 | 832 | 91.85 | 1.600 |
| 28/09/2018 | vir-1 | 3 | - | 1,843,192 | 865 | 1,746,153 | 1,134 | 480 | 15,291 | 884 | 94.74 | 0.878 |
| 03/10/2018 | vir-1 | 4 | - | 1,355,795 | 1436 | 1,271,511 | 1,128 | 487 | 11,522 | 802 | 93.78 | 1.467 |
| 19/10/2018 | VIRc | 3 | - | 1,678,723 | 1548 | 1,570,040 | 2,127 | 500 | 11,471 | 849 | 93.53 | 0.887 |

**Supplementary table 2.**
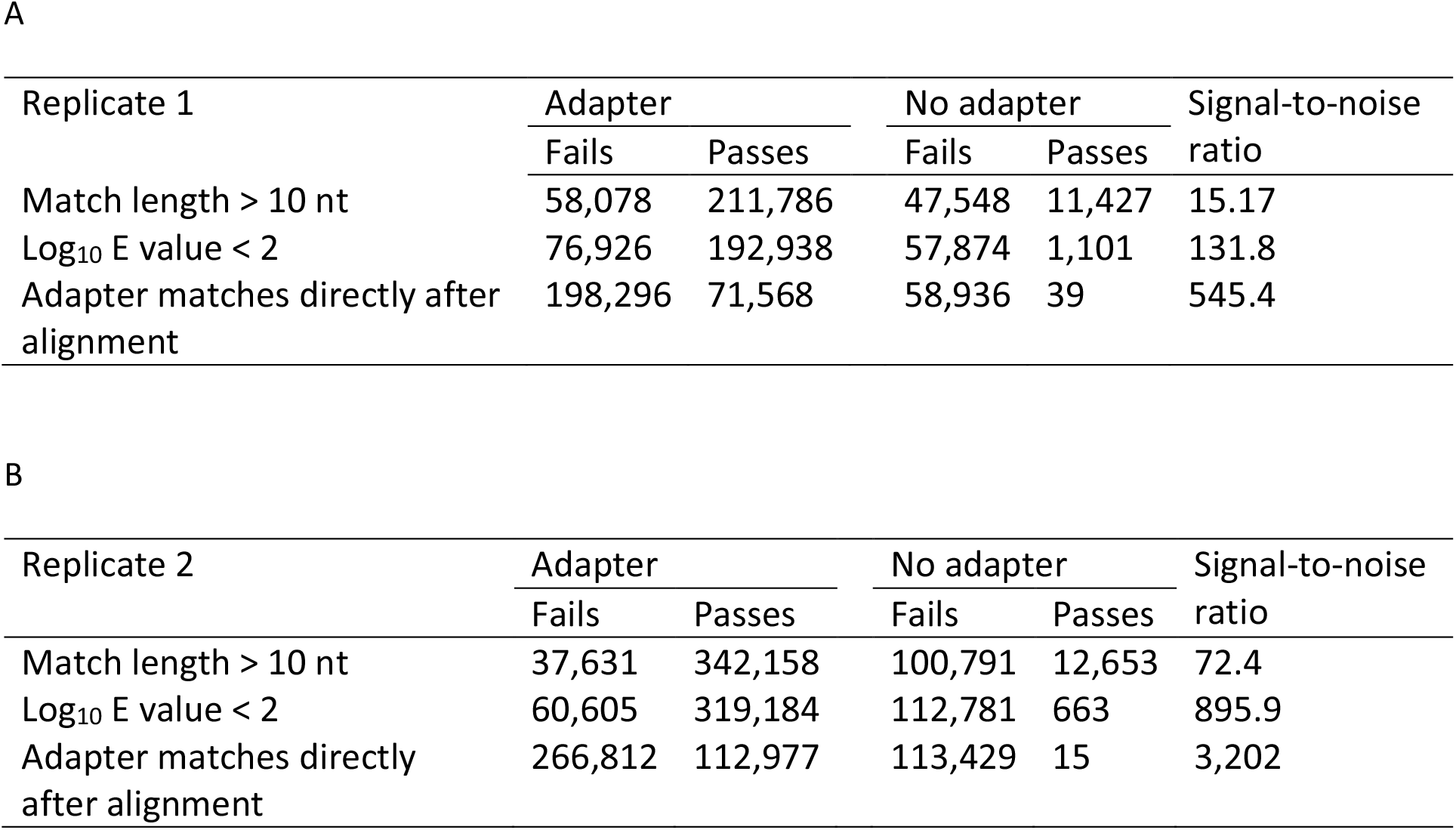
Adapter detection using BLASTN rules approach. The number of reads with adapters were detected in two biological replicates (Tables A and B respectively) of Col-0 sequenced with and without adapter ligation protocol. Rules are applied cumulatively (i.e. row one shows reads that pass the first rule, row two shows reads that pass the first and second rules, etc.). The signal-to-noise ratio shows the number of positive examples detected using rules in the adapter-ligated dataset divided by the number of false positives from the dataset collected without adapters. **[Linked to Figure 3].**

| Replicate 1 | Adapter |  | No adapter |  | Signal-to-noise ratio |
| --- | --- | --- | --- | --- | --- |
|  | Fails | Passes | Fails | Passes |  |
| Match length > 10 nt | 58,078 | 211,786 | 47,548 | 11,427 | 15.17 |
| Log<sub>10</sub> E value < 2 | 76,926 | 192,938 | 57,874 | 1,101 | 131.8 |
| Adapter matches directly after alignment | 198,296 | 71,568 | 58,936 | 39 | 545.4 |

**Supplementary table 2. Adapter detection using BLASTN rules approach.**
| Replicate 2 | Adapter |  | No adapter |  | Signal-to-noise ratio |
| --- | --- | --- | --- | --- | --- |
|  | Fails | Passes | Fails | Passes |  |
| Match length > 10 nt | 37,631 | 342,158 | 100,791 | 12,653 | 72.4 |
| Log<sub>10</sub> E value < 2 | 60,605 | 319,184 | 112,781 | 663 | 895.9 |
| Adapter matches directly after alignment | 266,812 | 112,977 | 113,429 | 15 | 3,202 |

**Supplementary table 3.**
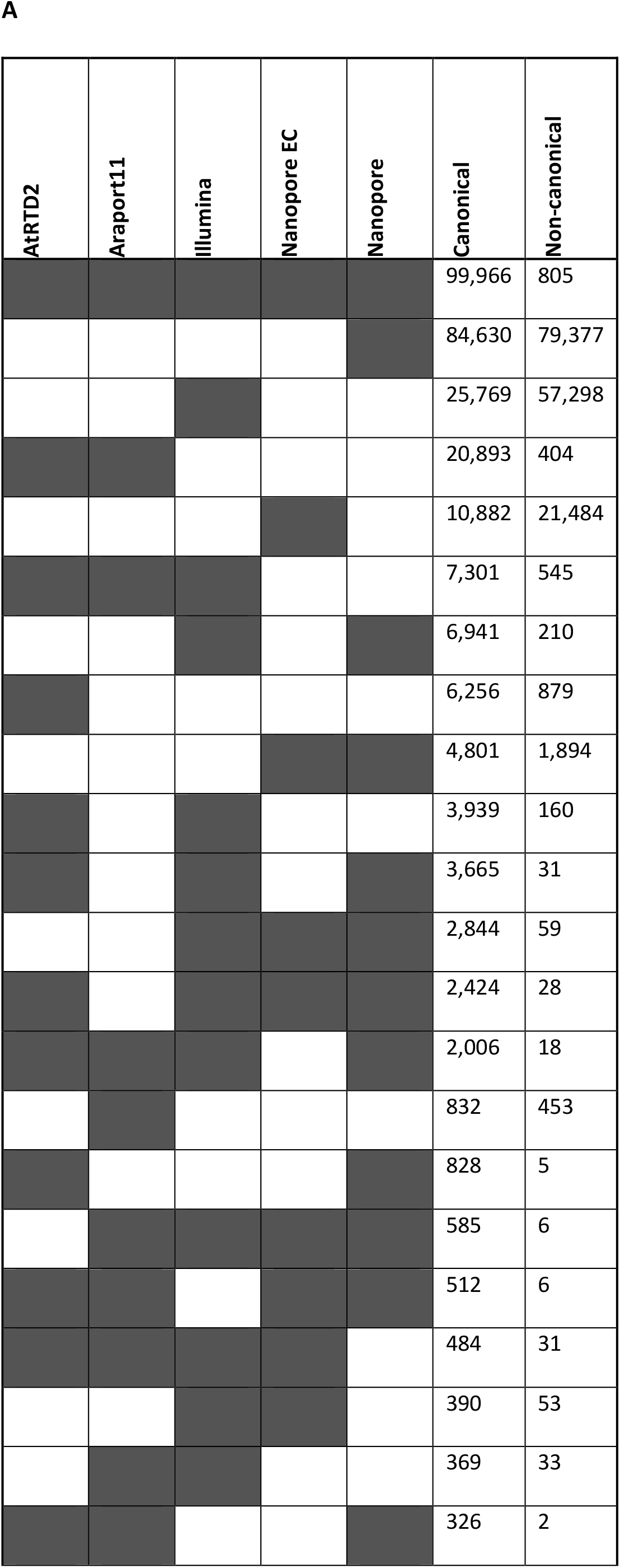
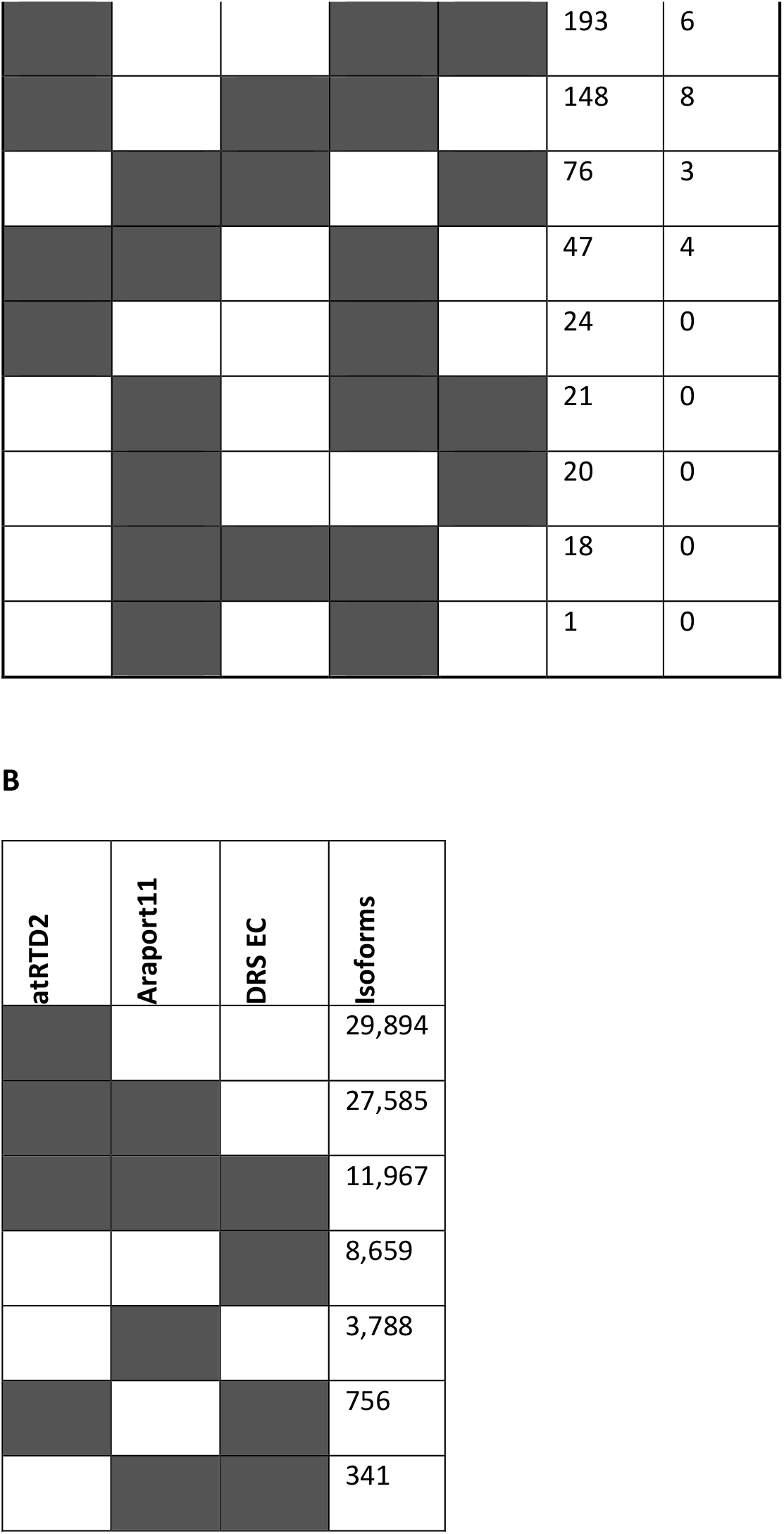
Splice junctions supported by nanopore DRS and Illumina RNAseq. Numbers are shown for (A) the unique splice junction set intersections upset plot (Figure 4B) and (B) unique linked splicing events upset plot (Figure 4C). Shaded cells denote sets included in the intersection for that row, while unshaded cells denote sets excluded from the intersection. Rows are sorted by the size of the intersection for canonical splice junctions. **[Linked to Figure 4].**

**Supplementary table 4.**
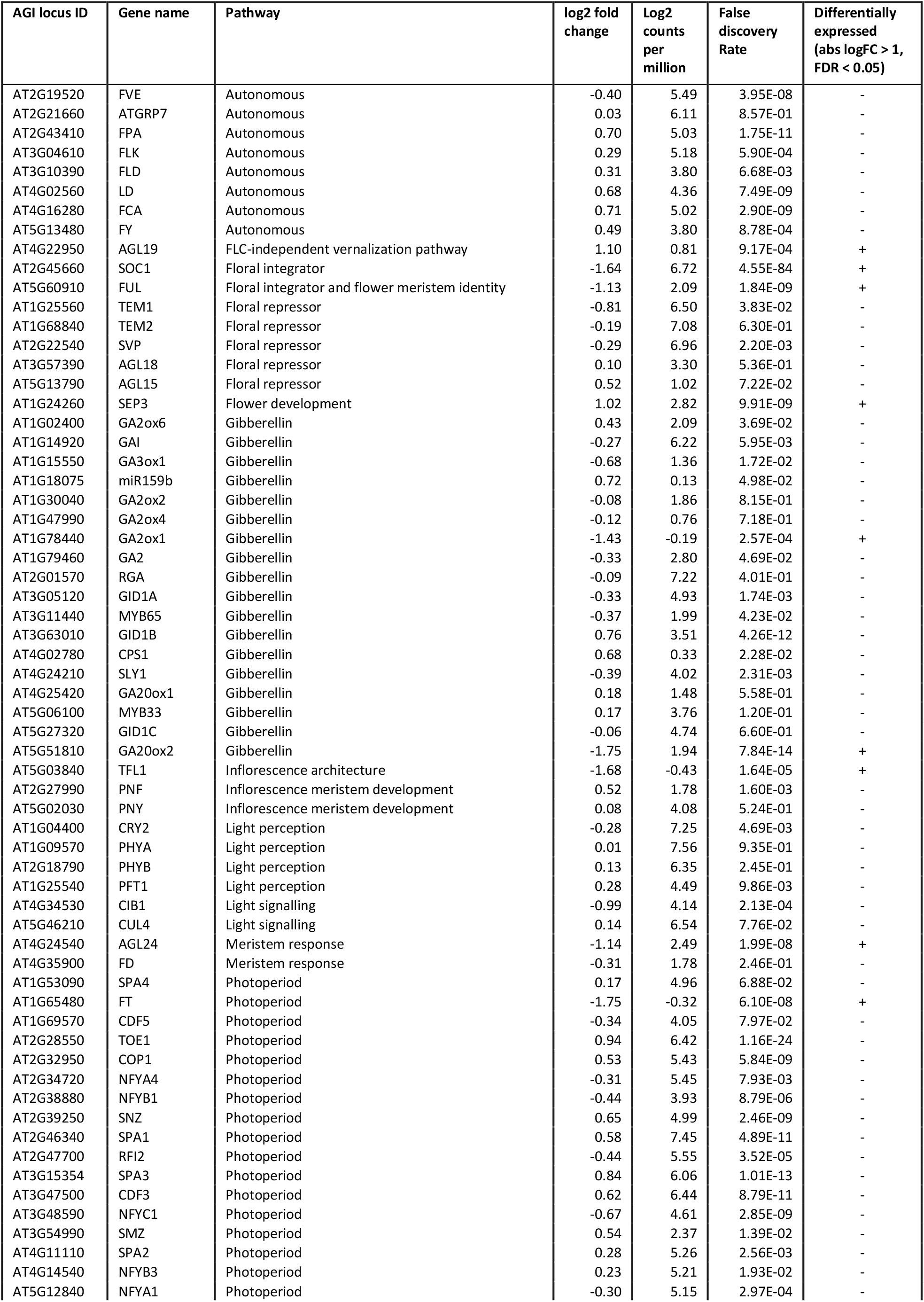
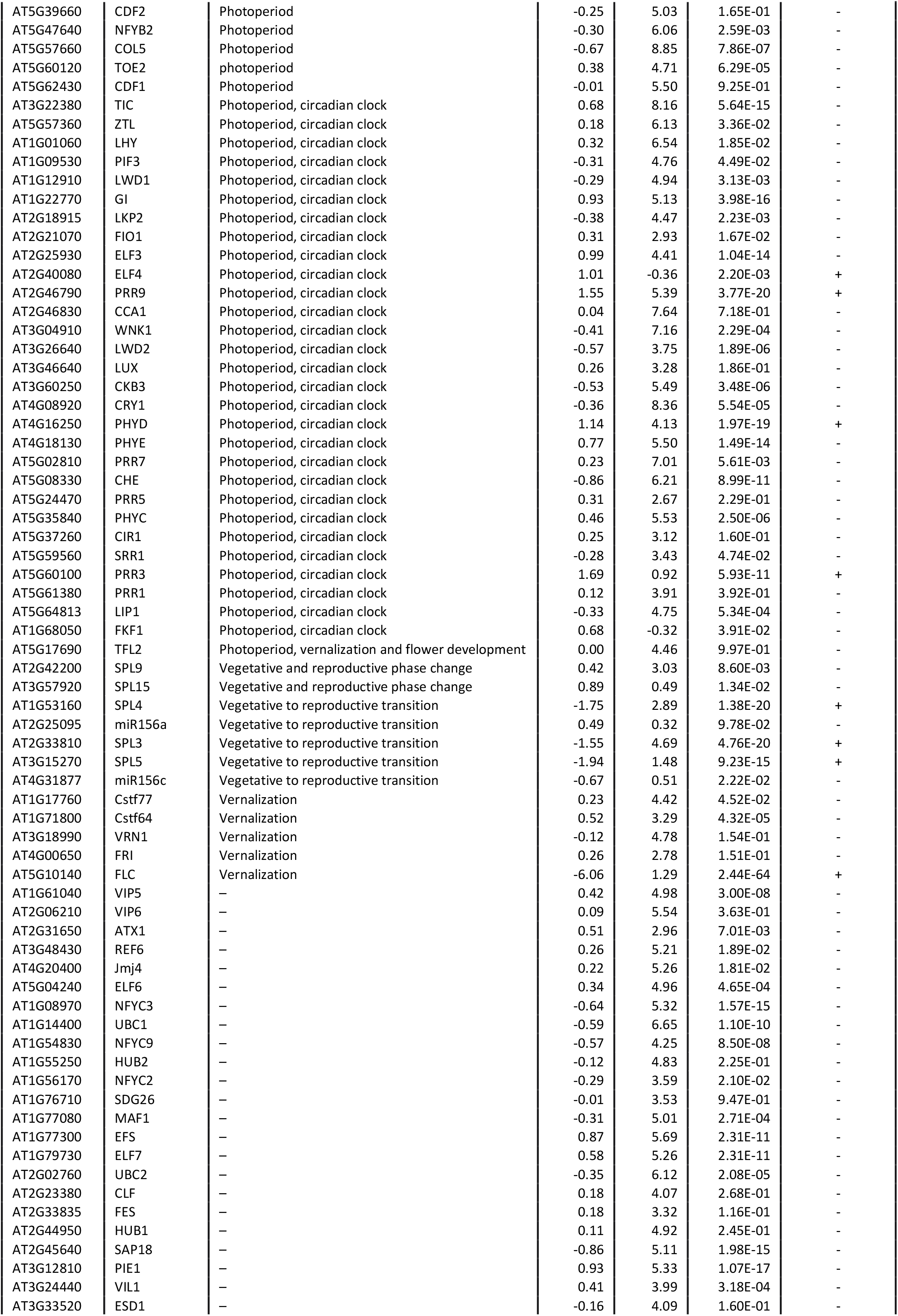
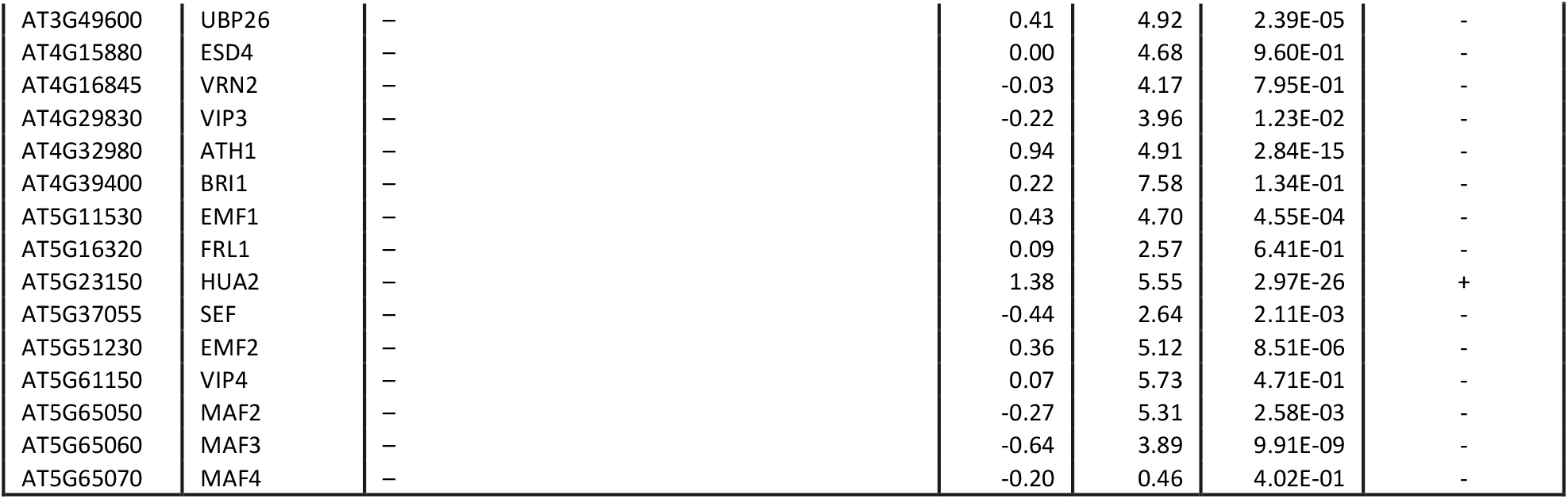
Flowering time gene expression. Change in gene expression of curated genes involved in flowering time in Arabidopsis, as detected using Illumina RNAseq for *vir-1* compared with the VIR-complemented line. In all, 12.2% of flowering time genes show a change in mRNA level expression in the *vir-1* mutant. Source of flowering time genes: George Coupland, Cologne: https://www.mpipz.mpg.de/14637/Arabidopsis_flowering_genes **[Linked to Figure 6].**

**Supplementary table 5.**
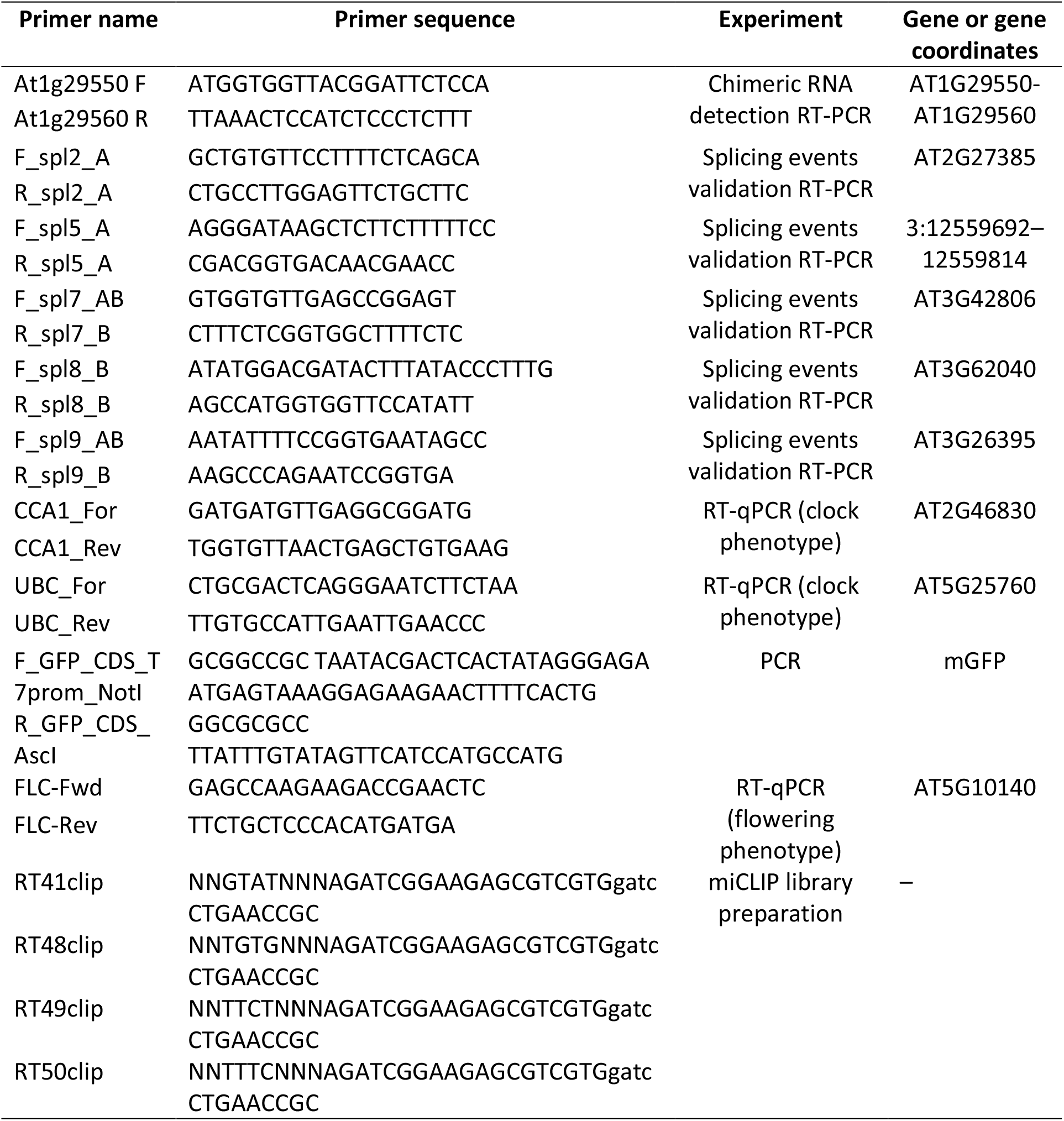
Primers used in this study.

